# Circuit-Based Understanding of Fine Spatial Scale Clustering of Orientation Tuning in Mouse Visual Cortex

**DOI:** 10.1101/2025.02.11.637768

**Authors:** Peijia Yu, Yuhan Yang, Olivia Gozel, Ian Oldenburg, Mario Dipoppa, L. Federico Rossi, Kenneth Miller, Hillel Adesnik, Na Ji, Brent Doiron

**Affiliations:** Department of Neurobiology, University of Chicago, Chicago, IL, USA; Grossman Center for Quantitative Biology and Human Behavior, University of Chicago, Chicago, IL, USA; Department of Physics, University of California, Berkeley, CA, USA; Helen Wills Neuroscience Institute, University of California, Berkeley, CA, USA; Department of Molecular and Cell Biology, University of California, Berkeley, CA USA; Department of Neuroscience and Cell Biology, Robert Wood Johnson Medical School, and Center for Advanced Biotechnology and Medicine, Rutgers University, Piscataway, NJ, USA; Department of Neurobiology, University of California, Los Angeles, CA, USA; Center for Theoretical Neuroscience, Department of Neuroscience, Columbia University, New York, New York, USA; Swartz Program in Theoretical Neuroscience, Columbia University, New York, New York, USA; Mortimer B. Zuckerman Mind Brain Behavior Institute, Columbia University, New York, New York, USA; Center for Neuroscience and Cognitive Systems, Italian Institute of Technology, Rovereto (TN), Italy; Kavli Institute for Brain Science, Columbia University, New York, New York, USA; Molecular Biophysics and Integrated Bioimaging Division, Lawrence Berkeley National Laboratory, Berkeley, CA, USA; Department of Statistics, University of Chicago, Chicago, IL, USA

## Abstract

In sensory cortex of brain it is often the case that neurons are spatially organized by their functional properties. A hallmark of primary visual cortex (V1) in higher mammals is a columnar functional map, where neurons tuned to different stimuli features are regularly organized in space. However, rodent visual cortex is at odds with this rule and lacks any spatially ordered functional architecture, and rather neuron feature preference is haphazardly organized in patterns termed ‘salt-and-pepper’. This sharp contrast in feature organization between the visual cortices of rodents and higher mammals has been a persistent mystery, fueled in part by abundant evidence of conserved cortical physiology between species. In this work, we applied a novel GCaMP indicator that are localized in the nucleus of neurons during two-photon imaging in mouse V1, which enabled us to overcome most spurious spatially correlated activity due to fluorescence contamination, and to ensure a faithful observation of functional organization over space. We found that the orientation tuning properties of distant neuron pairs (> 20 *µm*) are irregularly and randomly organized, while neuron pairs that are extremely close (< 20 *µm*) have strongly correlated orientation tuning, indicating a narrow yet strong spatially clustered organization of orientation preference, which we term ‘micro-clustered’ organization. Exploring a circuit-based model of recurrently coupled mouse V1 we derived two key predictions for the ‘micro-cluster’: spatially localized recurrent connections over a comparable narrow spatial scale, and common relative spatial spreads of balanced excitation and inhibition in the network over broad spatial scales. These predictions are validated by both anatomical and optogenetic-based physiological circuit mapping experiments. Altogether, our work takes an important step in building a circuit-based theory of visual processing in mouse V1 over spatial scales that are often ignored, yet contain powerful synaptic interactions.

## Introduction

A general dictum in systems neuroscience is that stimulus or behavioral tuning of a single neuron’s response is a reflection of the circuit structure within which the neuron is embedded. Prior experimental work on recurrent circuitry in the neocortex suggests that neuron pairs that are physically closer to one another have a higher likelihood of direct synaptic connections (Campagnola et al., 2022; Holmgren et al., 2003; Levy and Reyes, 2012; Rossi et al., 2020; Seeman et al., 2018). Thus, one may expect that the distribution of stimulus feature preference over a cortical population may show a strong spatial component, where neurons that are near one another will be similarly tuned. This is indeed the case in the columnar structure of orientation tuning in the visual systems of cat (Hubel and Wiesel, 1962), ferret (Chapman et al., 1996), tree shrew (Bosking et al., 1997) and several species of non-human primates (Espinosa and Stryker, 2012; Hubel and Wiesel, 1968); the tonotopic organization of auditory cortex (Schreiner and Winer, 2007); and the barrel structures in somatosensory whisker system (Andermann and Moore, 2006). However, while neurons in the primary cortex of the visual system of rodents (V1) show tuned responses to oriented gratings, calcium imaged responses show that the population famously lacks any clear spatial organization of tuning preference and rather has a ’salt-and-pepper’ organization (Kaschube, 2014; Ohki et al., 2005). This apparent disorder is in seeming opposition to the known spatial rules for synaptic connectivity in mouse V1 that mirror other cortices (Campagnola et al., 2022; Rossi et al., 2020), which could in principle impart some spatial organization to functional responses.

After the original reports of salt-and-pepper organization of feature preference in mouse V1 (Bonin et al., 2011; Ohki et al., 2005; Ohki and Reid, 2007), several studies have revisited the finding and presented conflicting results. Ringach et al., 2016 reported a spatially clustered organization of stimulus feature preference that spans approximately a hundred microns horizontally across L2/3 of mouse V1. In contrast, other imaging experiments in mouse V1 show either a much narrower scale of clustered organization (Kondo et al., 2016), or a complete lack of spatial organization (Han et al., 2019). This disagreement about a fundamental aspect of cortical representation is problematic, and is an obstacle to relating functional organization to the underlying circuit structure in mouse V1 – one of the best characterized neuronal regions.

To resolve this fundamental disagreement about the organization of orientation preference in mouse V1, in this study we leveraged nuclear localized GCaMP6 calcium indicators that eliminate a potential key technical challenge in previous studies: the ‘neuropil’ contamination of cellular signals by surrounding background tissue (Dipoppa et al., 2018; Gobel and Helmchen, 2007; Pachitariu et al., 2017). Since neuropil is spatially dispersed, it may produce spurious correlations in activity between nearby neurons, potentially masking the true functional organization. By restricting fluorescence to cell nuclei, our approach minimizes this confounding effect. Using this refined approach, we discovered a distinct organization of orientation preference in mouse V1: neurons form strong spatial clusters at an extremely fine scale (∼ 20 *µm*), which we term ’micro-clustered’ organization. This organization, observed in both layer 4 (L4) and layer 2/3 (L2/3), reconciles the apparent contradiction between earlier studies and suggests a precise spatial structure that may bridge the gap between cellular organization and underlying circuit connectivity.

A micro-clustered organization of orientation preference presents a new mechanistic puzzle: neurons cluster at a fine spatial scale (∼ 20 *µm*), yet typical recurrent connections in mouse V1 spread over much broader distances (100 ∼ 200 *µm*) (Campagnola et al., 2022; Holmgren et al., 2003; Levy and Reyes, 2012; Rossi et al., 2020; Seeman et al., 2018). We propose two potential resolutions to this apparent mismatch. First, there may exist an additional, previously underappreciated component of synaptic connectivity operating at fine spatial scales (5 ∼ 20 *µm*) (Li et al., 2012). Second, previous theoretical work on spatially extended networks suggests that the relationship between connectivity and functional organization is not necessarily direct - the balance between excitation (E) and inhibition (I) can decouple spatial patterns of activity from underlying circuit connectivity (Rosenbaum and Doiron, 2014; Rosenbaum et al., 2017). Building on these insights, we developed a circuit-based model of mouse V1 that combines two distinct organizational principles: a ‘macro-scale’ E/I balance that prevents spatial patterning at broader scales, and a ‘micro-scale’ E/I imbalance that enables the emergence of fine-scale clusters. This model makes specific, quantitative predictions about the spatial organization of feedforward (L4 to L2/3) and recurrent (L2/3 to L2/3) E/I connectivity required to produce the observed micro-clustered organization of orientation preference.

We tested these model predictions through multiple complementary experimental investigations. First, a reanalysis of anatomical pre-synaptic ensemble mapping with rabies tracing (Rossi et al., 2020) confirmed the conditions necessary for the absence of macro-scale spatial organization. Second, targeted circuit mapping using two-photon holographic optogenetics (Oldenburg et al., 2024) revealed strong synaptic connections within micro-scale domains. Finally, analysis of trial-to-trial population co-variability (noise correlations) demonstrated the predicted narrow spatial scale organization. Altogether, our combined experimental and theoretical investigations provide a comprehensive chain of evidence for a novel functional micro-cluster organization of orientation preference in mouse V1. This fine-scale principle of circuit and functional organization, though typically overlooked, may serve essential roles in visual information processing within the rodent cortex.

## Results

### Micro-clustered organization of orientation tuning preference in L2/3 and L4 of mouse V1

Using viral transduction we labeled V1 L2/3 neurons (both excitatory and inhibitory) in wildtype mice with the cytosolically expressed genetically encoded calcium indicator GCaMP6s (Chen et al., 2013). After habituating the mice to head fixation, we characterized the visually evoked calcium responses of these L2/3 neurons in awake mice. We presented a pseudorandom sequence of sinusoidal grating stimuli drifting in one of 12 motion directions, interleaved with gray screens (10 trials each stimulus; see Methods). For each mouse, we acquired data from multiple image planes parallel to the cortical surface, with a vertical (laminar) step size of 20 *µm*. After image registration, active GCaMP6s^+^ neurons were detected by the analysis software platform *Suite2p* (Pachitariu et al., 2017) and outlined as regions of interest (ROIs) (Figure 1A). The direction tuning curve for each neuron was calculated from its trial averaged Δ*F*(*t*)/*F*_0_ transient (upper panel, Figure 1B) in response to each drifting direction (lower panel, Figure 1B; see Methods for details).

For visually responsive neurons (max trial-averaged Δ*F*(*t*)/*F*_0_ > 0.1) with significantly different responses across the 12 directions (one-way ANOVA test *P* < 0.05), we fit their tuning curves with a bimodal Gaussian function. Neurons with well fit tuning curves were defined as orientation selective (OS) neurons (see Methods), and only OS neurons were further analyzed. Across all imaging depths, all field of views (FOVs) and all mice, 22.6% of L2/3 neurons (n = 2246/9936, 2 mice) were labeled OS. Preferred orientations of these OS neurons were distributed over the full range of stimulus orientations (upper panel, Figure 1C; Figure S1), and the shapes of the tuning curves were heterogeneous (middle panel, Figure 1C; Figure S1). To quantify neuronal tuning we used an global orientation selectivity index (gOSI, see Methods) where gOSI = 0 implies no selectivity, and gOSI = 1 implies perfect selectivity. The gOSI distribution over OS neurons was unimodal with a mean value near 0.5 (Figure 1C, lower panel; Figure S1), indicating robust orientation selectivity in mouse V1, matching past reports (Bonin et al., 2011; Kaschube, 2014; Ohki et al., 2005; A. Y. Tan et al., 2011; Van Hooser et al., 2005). Using the preferred orientations of OS neurons, we constructed color-coded spatial maps of preferred orientation for different horizontal image planes in L2/3 (Figure 1A; Figure S1). Upon visual inspection, neurons with the same preferred orientation (color) appeared to be randomly distributed in the horizontal plane without obvious spatial structure, i.e. a “salt-and-pepper” spatial organization (Kaschube, 2014; Ohki et al., 2005). However, as mentioned in the introduction, there is significant disagreement in the literature surrounding the lack of spatial organization (Jimenez et al., 2018; Kondo et al., 2016; Ringach et al., 2016), so we next aimed to better quantify the extent of any spatial organization of orientation selectivity.

We examined how the similarity of orientation tuning varied with the cortical distance between pairs of neurons (i.e., distance between the center of mass of their cell bodies). Two metrics for the tuning similarity between a pair of neurons were used: one was the correlation coefficient between their tuning curves (inset of Figure 1D), and the other was the difference between their preferred orientation angles (inset of Figure 1E). The correlation metric takes into account the entire shape of tuning curves, while the difference in preferred orientations considers just the locations of peak tuning. For the moment, we investigate the horizontal spatial structure (i.e. “salt-and-pepper” or clustered) rather than vertical organization (i.e. columnar). As such we only consider the distance measured along horizontal cortical plane and ignore any vertical distance between neurons. Because of this the reported distance between neuron pairs may be smaller than the diameter of a neuron’s soma (∼ 10 *µm*) since overlapping nearby neuron pairs may nevertheless be offset vertically.

**Figure 1:**
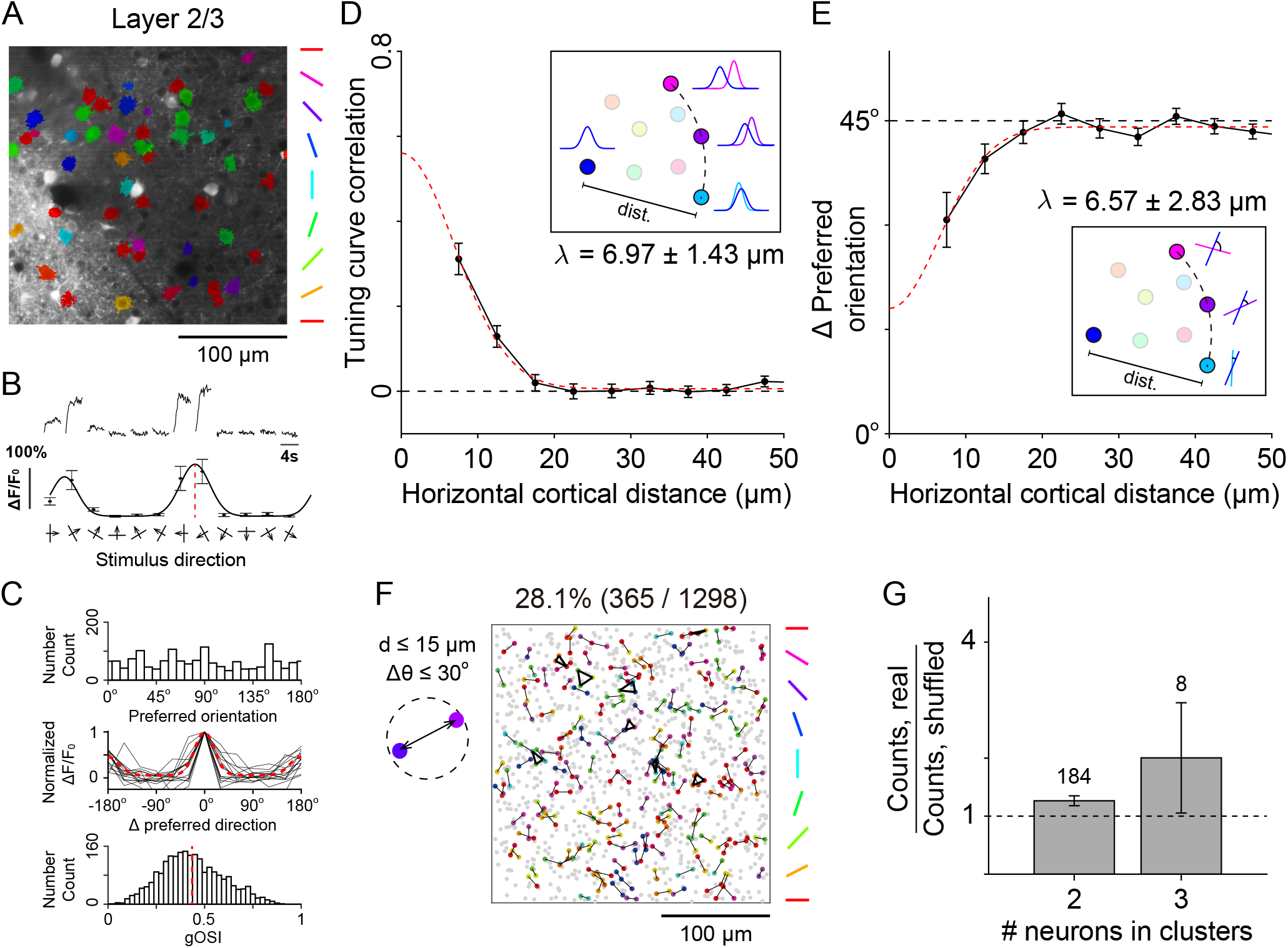
Micro-clustered spatial organization of orientation tuning over a fine spatial scale in L2/3 of mouse V1. (A) *In vivo* fluorescence image of an example optical plane of cytosolic GCaMP6^+^ L2/3 neurons. The detected regions of interest (ROIs) of orientation selective (OS) neurons are color coded by their preferred orientations. (B) The Δ*F*/*F*_0_(*t*) traces of transient calcium dynamics (top) as well as the tuning curve (bottom) of an example orientation selective (OS) neuron in L2/3, in response to drifting grating stimuli in 12 directions (average of 10 trials). Red dashed line marks the preferred direction. (C) Top: Distribution of the preferred orientation of all L2/3 OS neurons (n = 2246). Middle: Normalized tuning curves of 15 random individual examples (gray) as well as the population average (red dashed) of all L2/3 OS neurons, relative to preferred directions. Bottom: Distribution of the global orientation selectivity index (gOSI) of all L2/3 OS neurons. Red dashed line represents the mean value (0.435). (D) The correlation coefficient between tuning curves of pairs of neurons sharply decreases as a function of the horizontal cortical distance between neurons. This indicates a strong and significant orientation tuning similarity over a fine spatial scale, rather than a salt-and-pepper organization. Error bars represent standard error of the mean. The red dashed curve is the optimal fit to Gaussian function A·exp(−d^2^/2λ^2^)+b, where λ is the standard deviation / spatial width (±Δλ is the 95% confidence interval). Inset schematic: examples of tuning curve correlation between multiple pairs of neurons with the same horizontal cortical distance. (E) The difference between preferred orientation angles of neuron pairs sharply increases as a function of the horizontal cortical distance. Error bars, dashed curve and λ: the same as in (D). Inset cartoon: examples of difference between preferred orientation angles of multiple pairs of neurons with the same horizontal cortical distance. (F) L2/3 OS neurons on the same imaging plane that are close to one another (distance < 15 *µ*m) and have similar preferred orientations (difference < 30^*o*^) are interconnected and colored coded by their preferred orientations. Gray dots indicates neurons that are not members of a micro-cluster. Neurons from different imaging planes and different mice were vertically stacked. The proportion of neurons within micro-clusters is indicated above the panel. (G) The ratios of actual number counts of L2/3 micro-clusters (numbers above the bars) in different sizes (2: pair; 3: triplet) to that of randomly shuffled orientation maps. Error bars represent standard deviation of the mean.

A true salt-and-pepper organization predicts that both the tuning curve correlation and the difference between preferred orientation angles are statistically independent of cortical distance (dashed black line of zero correlation in Figure 1D, and 45^*o*^ angle difference in Figure 1E). However, our data shows that the tuning curve correlation starts from a high level for neuron pairs adjacent to each other (starting from 7.5 *µm* in Figure 1D), but sharply decays and asymptotes to zero when cortical distance is greater than ∼ 20 *µ*m (Figure 1D; Figure S2). In agreement with this, the difference between preferred orientation angles start from a low level for short cortical distances, but sharply increases and asymptotes to 45^*o*^ at longer distances (Figure 1E; Figure S2). Gaussian fits to the data yielded a spatial width (standard deviation) for orientation tuning similarity of ∼ 7 *µm* (dashed red lines in Figures 1D and 1E; spatial width λ reported in the panel). These results suggest that in L2/3 there is a strong and significant organization of tuning preference over a very fine spatial scale (essentially adjacent neurons), rather than a purely spatially disorganized tuning map. We label such a spatial organization as ‘micro-clustered’, where ‘micro’ represents a small spatial scale often overlooked in population analysis.

**Figure 2:**
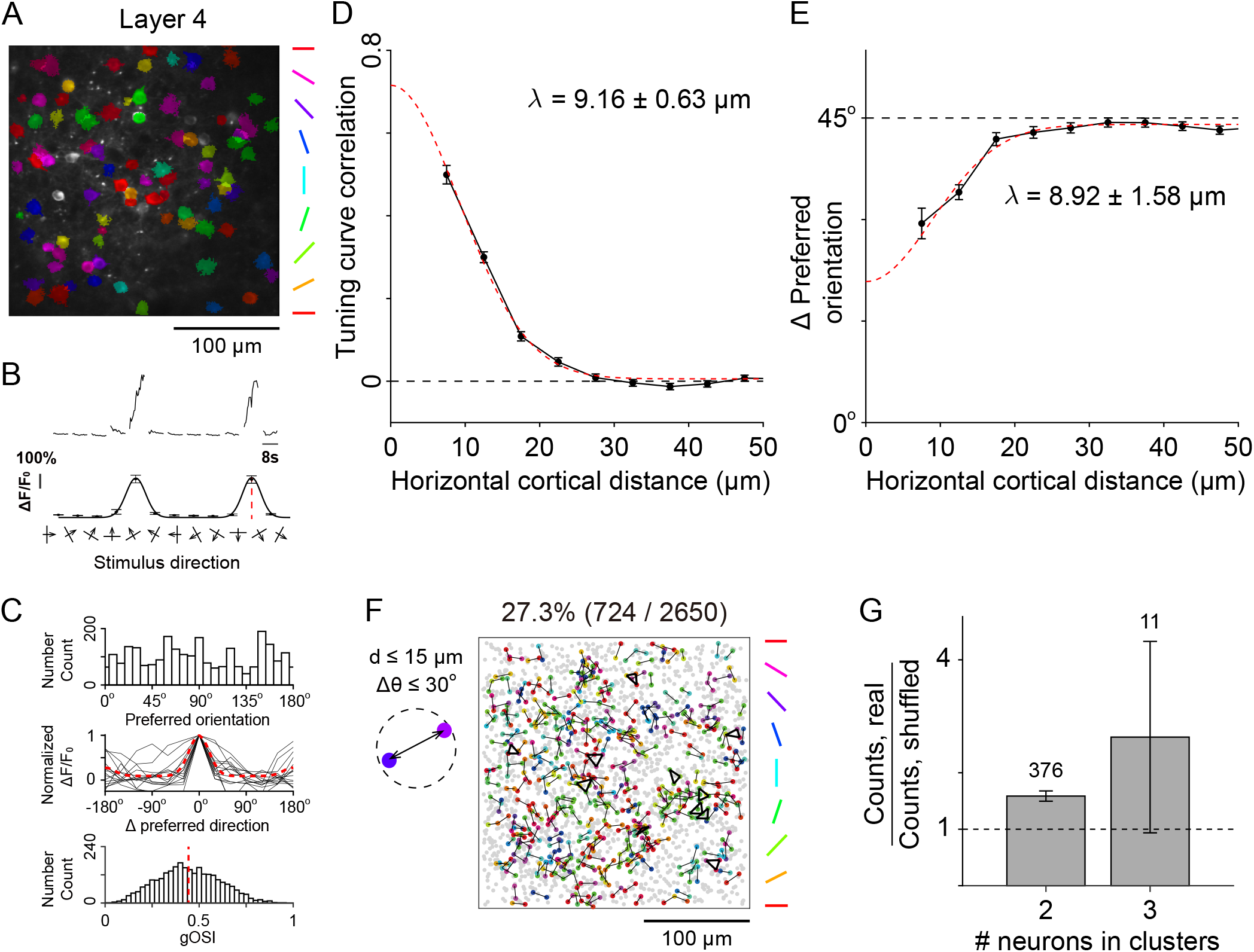
Micro-clustered spatial organization of orientation tuning over a fine spatial scale in L4 of mouse V1. The panels are organized as in Figure 1. (B) n = 2650 for L4 OS neurons; mean value of gOSI 0.443.

In the primary visual cortex a major source of input to a L2/3 neuron is from L4 (Douglas and Martin, 2004). We therefore asked whether the micro-clustered structure in L2/3 is inherited (in part) from a similarly micro-clustered organization in L4. We labeled V1 L4 excitatory neurons in *Scnn1a*-Tg3-Cre transgenic mice (Madisen et al., 2010) with cytosolically expressed GCaMP6s, and measured their calcium responses to drifting grating stimuli in head-fixed awake mice. Using the same image processing method and the same criteria of L2/3 OS neurons, we found 56.6% of L4 neurons (n = 2650/4684, 5 mice) were classified OS. The orientation tuning properties of L4 OS neurons were similar to that of L2/3 OS neurons (compare Figure 2A-2C to Figure 1A-1C). Importantly, L4 neuron pairs which are very close to one another on the same horizontal plane tend to have highly overlapping tuning curves (Figure 2D) and similar preferred orientation angles (Figure 2E), indicating a L4 micro-clustered organization similar to that in L2/3. While this suggests that the L2/3 clustering is inherited from L4, there is a complication: the spatial width of L4 micro-clusters is broader than that in L2/3 (9.16 ± 0.63 *µm* for L4 in Figure 2D and 2E, v.s. 6.97 ± 1.43 *µm* for L2/3 in Figure 1D and 1E). This cannot be explained by a simple feedforward propagation from L4 to L2/3, because any reasonable non-orientation selective axonal projection from L4 to L2/3 would always widen the pattern in the recipient layer (L2/3). Thus, a narrower micro-clustered organization in L2/3 than that in L4 suggests a non-trivial functional role of recurrent circuitry in L2/3.

To better quantify the extent of micro-clustered structure in the spatial maps of preferred orientation, we measured the proportion of neurons that were members of identified micro-clusters. For this purpose, we defined a micro-cluster to be a group of neurons in the same horizontal imaging plane with similar preferred orientations to each other (difference < 30^*o*^), and close to at least one other member (distance < 15 *µm*; schematic at top left of Figure 1F and 2F). In both L2/3 (nuclear targeted GCaMP recordings; see next chapter for details) and L4 we found that more than a quarter of OS neurons were within micro-clusters (interconnected colored dots in Figure 1F and 2F. L2/3: 28.1%; L4: 27.3%). Most of these micro-clusters contained just 2 neurons, while a few of them contained 3 neurons (triplet) (numbers above the bars in Figure 1G and 2G). The number of micro-clustered pairs or triplets was higher than that of randomly shuffled orientation maps (the ratio levels in Figure 1G and 2G).

Apart from measuring the horizontal (within the imaging plane) extent of micro-clusters, we also investigated the vertical distribution of spatial organization of L2/3 and L4 orientation tuning. We considered the tuning curve correlation as well as the difference between preferred orientation angles of neuron pairs whose horizontal cortical distance was less than 7.5 *µm*. The vertical cortical distance between a pair of neurons was defined as the distance between the depth of their optical imaging planes. We observed a high tuning similarity (high tuning curve correlation, Figure S3 left; and similar preferred orientations, Figure S3 right) for neuron pairs within a short vertical distance to one another in both L2/3 (λ = 14.7 *µm*) and L4 (λ = 15.6 *µm*). This indicates a non-columnar, locally clustered functional organization spanning tens of microns vertically within mouse V1. Taken together, our analysis supports a discernible micro-clustered organization of orientation tuning as compared to a randomized “salt-and-pepper” map in both L2/3 and L4 of mouse V1.

**Figure 3:**
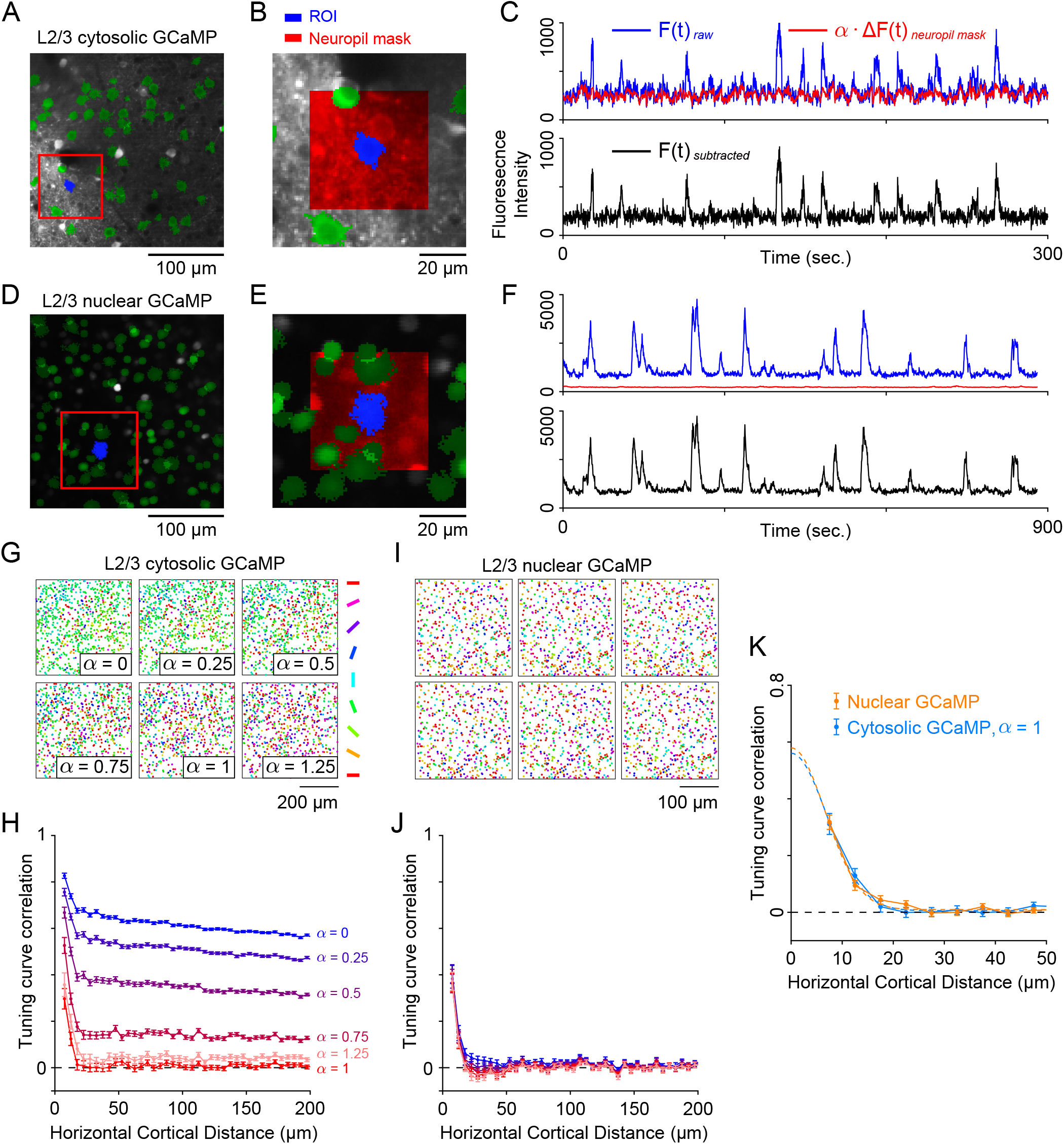
Measuring the effect of neuropil subtraction on micro-clustered organization. (A) The same example fluorescence image of cytosolic GCaMP^+^ L2/3 OS neurons as Figure 1C. The red square region is magnified in (B). (B) The ROI of an example cytosolic GCaMP^+^ L2/3 neuron (blue), as well as its neuropil mask (red), defined as the surrounding square region excluding any other ROIs (green). We suppose that the out-of-focus neuropil contamination within the ROI is proportional to the average signal in the corresponding neuropil mask, weighted by the correction scaling factor *α*. (C) *F*(*t*) traces of the raw ROI signal (blue), the estimated neuropil contamination signal (red, with *α* = 1), as well as the estimated neuronal response after subtraction (the bottom panel), for the example ROI in (B). (D-F) Analysis similar to (A-C) for an example nuclear-targeted GCaMP6^+^ L2/3 neuron. The neuropil contamination is largely reduced compared with cytosolic GCaMP^+^ neuron (compare panels C and F). (G) The color-coded spatial maps of preferred orientations for cytosolic GCaMP^+^ L2/3 OS neurons varies greatly with different values of *α*. Neurons from different imaging planes and different mice were stacked. (H) The correlation coefficient between tuning curves of cytosolic GCaMP^+^ L2/3 OS neurons as a function of cortical distance also varies greatly with different values of *α* (color-coded). (I, J) Analysis similar to (G, H) for nuclear-targeted GCaMP6^+^ L2/3 OS neurons, while both the orientation maps and the distance dependent functions of tuning curve correlation were almost independent of *α*. (K) The distance dependent function of tuning curve correlation for nuclear-targeted GCaMP6^+^ L2/3 neurons (orange) approximately matches the case of *α* = 1 for cytosolic GCaMP dataset (blue). Error bars: data; dashed line: Gaussian fit.

### Correcting for spurious micro-clusters due to neuropil contamination

One concerning issue for two-photon calcium imaging using cytosolic expression of GCaMP is that the recorded signal assigned to a particular ROI could be contaminated by a background neuropil signal. This neuropil signal originates from activity-induced fluorescence of dendrites, axons and glia surrounding the ROI (Pachitariu et al., 2017). This is of particular concern for our study, since the neuropil contamination signal is spatially diffuse and could create spurious spatial correlations of neuronal responses, especially over narrow spatial scales.

To correct for any neuropil contamination, we adopted past methods to subtract an estimate of the neuropil signal from the raw ROI signal (Dipoppa et al., 2018). A neuropil mask was defined as the square region 30 *µm* away from the center of each ROI, excluding pixels of other neighboring ROIs located on the same image plane (L2/3 example in Figure 3A and 3B, L4 example in Figure S4A and S4B). We then assumed that the neuropil contamination within the ROI was proportional to the average fluorescence signal in the neuropil mask, weighted by a scaling coefficient *α*. The true calcium trace *F*(*t*) was estimated by subtracting this neuropil signal from the raw signal measured from the ROI, i.e. *F*(*t*) = *F*(*t*)_raw_ − *α* · Δ*F*(*t*)_neuropil mask_ (examples in Figure 3C, 3F and S4C; more details in Methods). We adopted the simplification that *α* is uniform for all cells. The optimal linear scaling coefficient *α* provides a residual signal that is the best measure of the true neuronal response. In practice such an optimal *α* is either estimated manually (Chen et al., 2013; Peron et al., 2015; Kondo et al., 2016) or through automated algorithms (Pachitariu et al., 2017; Dipoppa et al., 2018). Before determining the optimal *α* for our datasets, we first explored how our finding of tuning similarity over a fine spatial scale is affected by different values of *α*.

For small spatial distances between L2/3 neuron pairs there were significant tuning curve correlations present for all values of *α* (Figure 3H, cortical distances less than ∼ 20 *µm*). However, for small *α* there was an over-representation of spatially broad clusters of neurons with similar orientation preference (group of green dots in orientation maps when *α* = 0, 0.25 and 0.5, Figure 3G), as well as high tuning curve correlations over a long range of pairwise distances over hundreds of microns (high asymptotic values of curves when *α* = 0, 0.25 and 0.5, Figure 3H). By contrast, larger *α* values reduce the over-represented preferred orientations from the spatial map (*α* = 0.75, 1 and 1.25 in Figure 3G), and the tuning curve correlations asymptote to lower values at long distances (curves of *α* = 0.75, 1 and 1.25 in Figure 3H). In particular, in this example case *α* = 1 (the red curve in Figure 3H) shows a near-zero asymptotic value of tuning curve correlation.

When neuropil contamination was not corrected in L4, the tuning curve correlation did not asymptote to as high a value as that in L2/3 (compare Figure S4E and Figure 3H). This was likely caused by the sparser expression of cytosolic GCaMP in L4 (i.e. only in Scnn1a-expressing excitatory neurons in L4 near the virus injection sites) than that in L2/3 (i.e., all neurons, from L1 to L5, both excitatory and inhibitory, near the virus injection sites), and thus a much reduced neuropil signal. Similarly, the asymptotic value of tuning curve correlation in L4 goes to approximately 0 when *α* = 1 (the red curve in Figure S4E).

These L2/3 and L4 results raise the concern that cytosolic GCaMP imaging measurements of long-range distance tuning similarity observed in awake mouse V1 might be artifactual, due to improper neuropil subtraction owing to a sub-optimal choice of *α* (Figure 3H). Thus, extreme care must be used in choosing an appropriate *α* to remove any spurious spatial structure of orientation tuning. In the following section we use nuclear-targeted GCaMP to eliminate fluorescence expression in the neuropil and provide a ground truth dataset so as to calibrate *α* directly.

### Nuclear-targeted GCaMP provides ground truth response and guideline for optimal neuropil subtraction

Nuclear localized versions of GCaMP6 (AAV-H2B-GCaMP6s) restrict fluorescence signals to the nucleus of the cell, so neuropil fluorescence from the surrounding region of an ROI is largely absent (compare Figures 3A and 3D). While nuclear-targeted GCaMP has activation and decay time constants that are much slower than that of cytosolic GCaMP, it can reliably report the tuning properties of neurons (Barchini et al., 2018).

We labeled L2/3 pyramidal neurons in mouse V1 with nuclear-targeted GCaMP and performed stimuli presentation, imaging, ROI identification, analysis of tuning curves and orientation maps identical to that performed for the cytosolic GCaMP analysis (Figure 3D-3E). We also applied the same neuropil subtraction method with nuclear-targeted GCaMP measurements (Figure 3F). In contrast to the cytosolic GCaMP data (example in Figure 3C), there was no neuropil contamination in nuclear-targeted GCaMP labeled cells (example in Figure 3F). The preferred orientation maps, and the tuning curve correlation as a function of cortical distance, were also independent of the neuropil subtraction scaling factor *α* (Figure 3I and 3J). Critically, the tuning curve correlations asymptote to near zero for distances of 100 ∼ 200 *µm* (Figure 3J), which were a near quantitative match to the case of *α* = 1 for cytosolic GCaMP measurements (compare blue and orange curves in Figure 3K). Thus, *α* = 1 is approximately the scaling factor which best removes neuropil activity and optimally reveal neuronal responses for our L2/3 cytosolic GCaMP dataset. These results suggest that nuclear-targeted GCaMP measurement can provide a ground truth to guide neuropil subtraction for cytosolic GCaMP measurements. The agreement between these two distinct datasets give strong support to a micro-clustered organization of orientation tuning preference in L2/3 in awake mouse V1.

In L4 we do not have the reagent to specifically express nuclear-targeted GCaMP. Thus, we chose *α* = 1 as our estimate of the optimal *α*, where the tuning curve correlation asymptotes to near zero for long distances (Figure S4E), in accordance with the L2/3 results. Finally, we note that for the initially presented cytosolic GCaMP L2/3 and L4 datasets (Figures 1 and 2), *α* = 1 was used to subtract the measured neuropil. We remark that in other datasets, the optimal *α* may differ from 1. Rather, the correct choice of *α* may depend on the varied conditions under which a dataset is imaged. For example, enlargement along the axial direction of the excitation focus, caused by degradation of image quality by the brain or imaging with a lower numeric aperture microscope objective, would lead to proportionally larger contamination from the neuropil (Ji et al., 2012; Wang et al., 2014).

**Figure 4:**
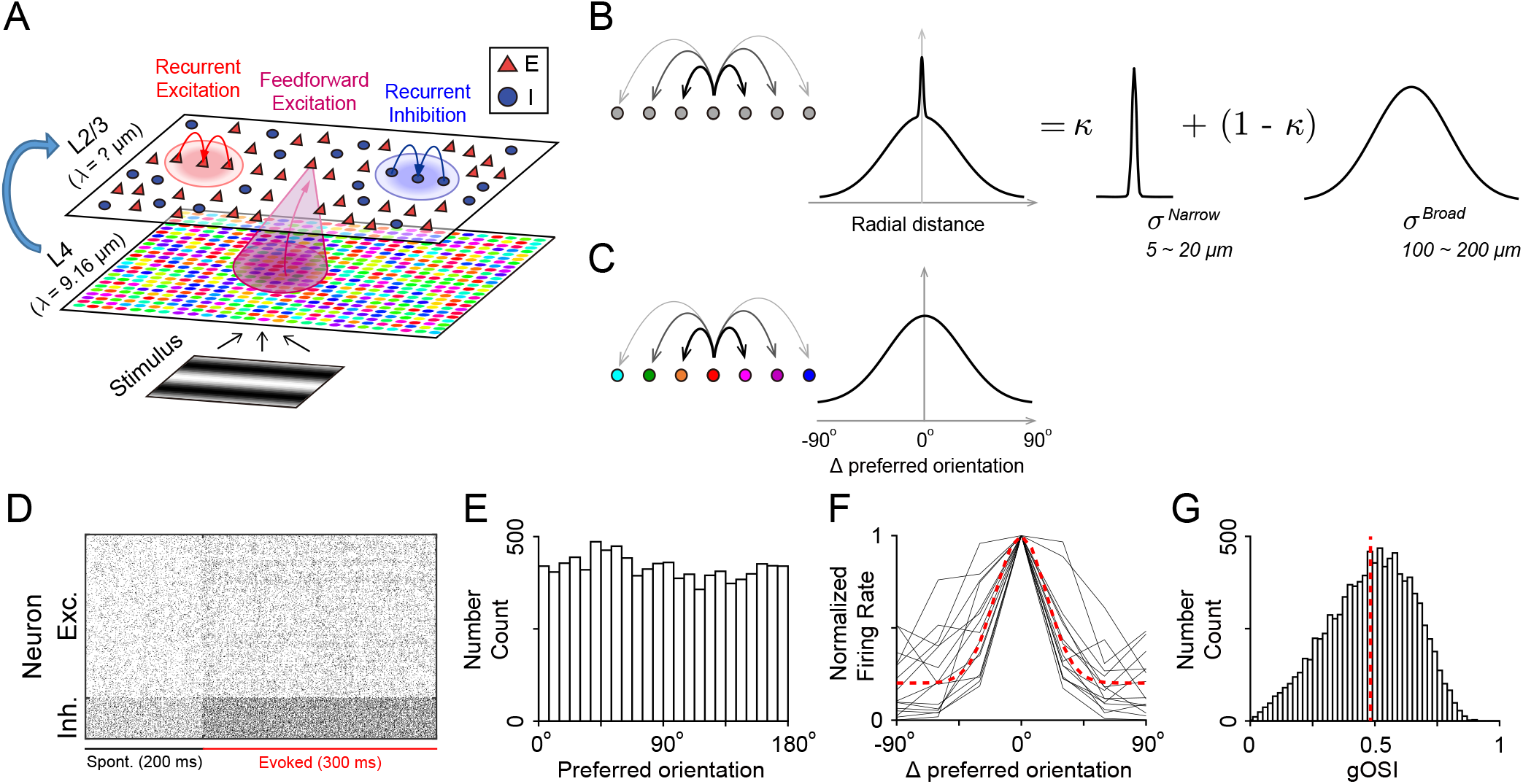
The framework of L4 – L2/3 spiking network model with broad and narrow connectivity. (A) Schematic of the two-layer network model. All neurons are arranged on two-dimensional squared grids. L4 contains Poisson-spiking neurons with Gabor receptive fields, receiving oriented grating visual inputs. The preferred orientations of L4, represented by different colors, are assigned according to the distance dependent function of tuning curve correlation in the data. L2/3 contains excitatory (E, red triangles) and inhibitory (I, blue circles) exponential integrate-and-fire neurons, receiving feedforward projections from L4 and recurrent projections from other L2/3 E and I neurons (magenta, red and blue footprints respectively). (B) The probability of feedforward and recurrent connections between two neurons decay with the radial distance between them, modeled as a sum-of-Gaussians-shaped function. The narrower Gaussian component, with ratio κ, has a spatial width (standard deviation) *σ*^Narrow^ of 5 ∼ 20 *µm* (similar to the width of micro-clusters). The broad component, with ratio 1 − *κ*, has a spatial width *σ*^Broad^ about 100 ∼ 200 *µm*. (C) The probability of feedforward and recurrent connections also depends on the difference between preferred orientations passed through a Gaussian-shape function. (D) Example of spike rasters of 10000 L2/3 E neurons and 2500 L2/3 I neurons during a 200 ms spontaneous activity and a 300 ms evoked activity. (E) Histogram distribution of the preferred orientations of all L2/3 E neurons. (F) Normalized tuning curves of 15 random individual examples (gray) as well as the population average (red dashed) of all L2/3 E neurons, relative to preferred orientations. (G) Histogram distribution of global orientation selectivity index (gOSI) of all L2/3 E neurons. Red dashed line marks the mean value.

### Broad and narrow connectivity in a network model of L4 and L2/3 of mouse V1

The strong experimental evidence of a micro-clustered organization of orientation tuning preference in both L2/3 and L4 populations prompted the following question: what circuit structures can support such a fine spatial scale functional organization? To begin to answer this question we constructed a two-layer network of model neurons arranged on two-dimensional lattices, modeling L4 and L2/3 of mouse V1 respectively (Figure 4A, see Methods). We focused our modeling efforts on determining the circuit requirements for the L2/3 network to inherit and sharpen the micro-clustered organization from L4, and we leave the genesis of L4 micro-clustering to a future study.

Our model contains 10000 L4 orientation-tuned Poisson-spiking neurons with preferred orientation angles that are spatially organized with the same distance dependent function of tuning curve correlation as in the data (Figure 4A, colored dots on the L4 sheet). This micro-clustered organization in L4 is enforced, and as such we omit any recurrent wiring within L4. The L2/3 network contains *N*_*E*_ = 10000 excitatory (E) and *N*_*I*_ = 2500 inhibitory (I) exponential integrate-and-fire model neurons (Gerstner et al., 2014; see Methods), each receiving feedforward excitatory synaptic inputs from L4 neurons, as well as recurrent synaptic excitation and inhibition from other L2/3 E and I neurons (Figure 4A, magenta, red and blue footprints on the L2/3 sheet respectively). For these three types of connections, the probability of a connection between two neurons decreases with the distance between the neuron pair. Past anatomical and electrophysiological studies of cortical circuits report spatial scales of synaptic wiring that span 100 ∼ 200 *µm* (Campagnola et al., 2022; Holmgren et al., 2003; Levy and Reyes, 2012; Rossi et al., 2020; Seeman et al., 2018). We label these scales as *broad* relative to the fine spatial scale of micro-clustered organization. The central model assumption of our study involves hypothesizing a second, *narrow* scale of synaptic wiring that is comparable to the 5 ∼ 20 *µm* scale of micro-clustered organization. Specifically, we postulate the following form for the connection probability between a pair of neurons separated by radial distance *d* (Figure 4B):

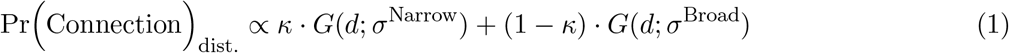

The function *G*(*d*; *σ*) is a normalized two-dimensional Gaussian-shaped function exp(−*d*^2^/2*σ*^2^)/(2*πσ*^2^), where *σ* is the spatial width (standard deviation) of synaptic coupling. In our model *d* is measured along the two-dimensional plane for L2/3 → L2/3 recurrent connections, and measured parallel to the plane for L4 → L2/3 feedforward connections. Eq. (1) decomposes the spatial profile of connection probability into a linear combination of two Gaussian functions, one with a broad spatial scale (characterized by *σ*^Broad^) and one with a narrow spatial scale (*σ*^Narrow^). The parameter *κ* is the ratio of the narrow component to the full connectivity. This sum of Gaussians connection probability indicates that neighboring neuron pairs tend to have much stronger effective connections than that of a single Gaussian-shaped function (the sharp peak in the center, Figure 4B). To cover the spatial scale of *σ*^Narrow^, we set the interval between neighboring L4, L2/3 E and L2/3 I neurons in the grid as 7.5 *µm*, 7.5 *µm* and 15 *µm*, respectively.

Apart from connectivity depending on the spatial distance between a neuron pair, we also assume that neuron pairs that prefer similar stimulus orientations are more likely to be connected, i.e. a ‘like-to-like’ feature-dependent rule that is known to exist in mouse V1 (Cossell et al., 2015; Ko et al., 2013; Rossi et al., 2020). These connections are also probabilistic and obey a Gaussian-shaped function (Figure 4C):

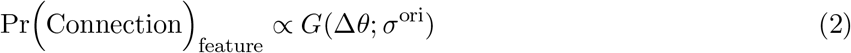

where Δ*θ* is the difference between preferred orientation angles of neurons, and *σ*^ori^ is the width along the orientation axis. Finally, we assume that the joint probability distribution for connectivity is the product of the spatial dependent term and the feature dependent term, i.e. these two factors are statistically independent:

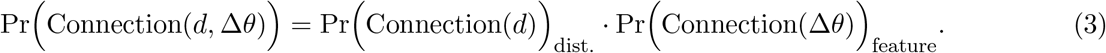

With standard parameters (see Methods) the network model produced irregular spiking activity in L2/3, that was roughly asynchronous in both spontaneous and evoked stimulus conditions (Figure 4D). Similar to the L2/3 data (Figure 1B), the L2/3 network model showed approximately uniform distribution of preferred orientation values (Figure 4E), heterogeneous but sharp tuning curves (Figure 4F), and robust orientation selectivity (quantified by gOSI, Figure 4G). If we omit the feature-dependent component of synaptic wiring, neurons continue to show orientation selectivity (Figure S5), as in previous studies of balanced networks (Hansel and van Vreeswijk, 2012; Pattadkal et al., 2018; Pehlevan and Sompolinsky, 2014). However, with our model parameters without feature dependent wiring the gOSI distribution is noticeably lower, indicating an overall weaker neuronal selectivity (Figure S5). Because of this, in what follows we always include the feature dependent component of synaptic wiring. However, we will restrict our analysis to the spatial structure of cortical wiring. In the following section, we will use our L4 → L2/3 network model to explore how the three key connectivity parameters: (*σ*^Broad^, *σ*^Narrow^, *κ*) determine the spatial organization of L2/3 orientation preferences. We remark that since there are three projection classes: L4 → L2/3 (feedforward), L2/3 E → L2/3 (recurrent excitation) and L2/3 I → L2/3 (recurrent inhibition) then there are nine parameters in total that are of interest.

### Conditions on pattern-forming and non-pattern-forming regimes in L2/3

In this section we provide a qualitative sketch of our theory underlying how L4 spatial patterns are, or are not, inherited and shaped by L2/3 circuitry. Our work is based on well established theories about spatial pattern formation in recurrently coupled networks (Ermentrout, 1998; Murray, 2007), and takes inspiration from the application of these theories to the visual system of higher mammals (cats and primates) (Bressloff and Cowan, 2003; Kang et al., 2003).

We begin with an (arbitrary) reference neuron in L2/3 and consider the spatial organization of the firing rates of the remaining L2/3 population. Specifically, we measure the average firing rate deviations of neurons at a distance *d* relative to the reference neuron, labelled Δ*r*_L2*/*3_(*d*). Presenting a theory for firing rate deviations is simpler than one for the spatial profile of tuning correlations; however, correlations can nevertheless be determined by Δ*r*_L2*/*3_(*d*) (see the Supplementary Methods S1 for details). Under a linear approximation of neuronal transfer, Δ*r*_L2*/*3_(*d*) is proportional to the total difference of synaptic input currents, which are decomposed into the sum of feedforward and recurrent terms:

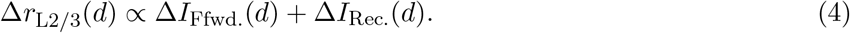

From Eq. (4) we will derive conditions for the spatial connectivity profiles of feedforward and recurrent wiring required to account for the presence or absence of spatial patterns in L2/3 activity.

The feedforward input current Δ*I*_Ffwd._(*d*) is generated from L4 activity which is then propagated to L2/3 via feedforward synaptic projections. We set the spatial organization of L4 activity, Δ*r*_L4_(*d*), to be a Gaussian with standard deviation λ_L4_, given by the Gaussian fits of the recorded L4 population response (Figure 2D; see the Supplementary Note for detail). For ease of analysis, we consider L4 → L2/3 projections to be translationally invariant, with a Gaussian shape and standard deviation *σ*_Ffwd_. The convolution of these two spatial distributions gives the effective spatial variance of the feedforward input Δ*I*_Ffwd_(*d*) as:

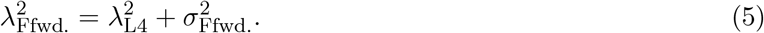

We note that in our framework the anatomical parameters for axonal projections are denoted by *σ*, and the effective spatial scales of response organizations are denoted by λ. In a similar fashion, the recurrent synaptic inputs Δ*I*_Rec._(*d*) are determined by L2/3 activity Δ*r*_L2*/*3_(*d*) and recurrent projection kernel 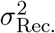. Their combination yields a spatial variance of:

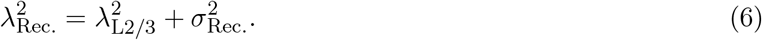

In the L4 – L2/3 network, the feedforward L4 → L2/3 synaptic input is excitatory (Figure 5A, the red Gaussian footprint), while the recurrent L2/3 → L2/3 input is inhibition dominated, which is true for awake mouse V1 (Haider et al., 2013; Sanzeni et al., 2020). Because of this we consider Δ*I*_Rec._(*d*) as a negative term (Figure 5A, the blue Gaussian footprint). A comparison between the spatial scales of feedforward excitation 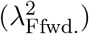 and (net) recurrent inhibition 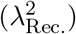 will give us mathematical conditions for whether spatial patterns will (or will not) occur in L2/3 activity.

We first make an ansatz (an educated mathematical assumption that must not lead to a contradiction) that the spatial pattern in the feedforward input is canceled by the recurrent pathway at all spatial locations. This will ensure that L2/3 will not exhibit spatially patterned activity (Rosenbaum and Doiron, 2014; Rosenbaum et al., 2017). Such cancellation occurs only when the spatial variances of the feedforward and recurrent inputs are the same: 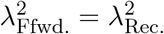. Equating the right hand sides of Eqs. (5) and (6) yields the following expression for the unknown space constant 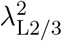:

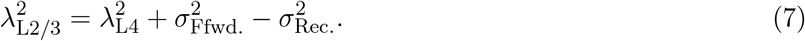

The solution Δ*r*_L2*/*3_(*d*) only exists if 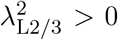 (because we can reject solutions with a zero or negative spatial scale). Thus, for the absence of any spatial pattern in L2/3 (as required by our ansatz), Eq. (7) gives the circuit constraint (Figure 5B_1_):

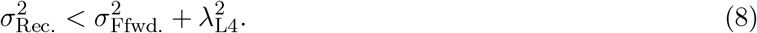

In other words, recurrent connections must be narrower than then spatial scale of the feedforward input in order for L2/3 to cancel any L4 patterning.

In contrast, when the recurrent projection width is so broad that we have

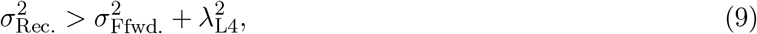

then our ansatz of input cancellation 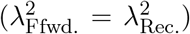leads to an impossibility, namely that 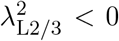. The conclusion must be that our cancellation ansatz over all spatial locations breaks down, and Eq. (4) cannot support a Δ*r*_L2*/*3_(*d*) solution that lacks a spatial pattern. In our case, the broad lateral inhibition from recurrent input will not counterbalance the narrower feedforward input, but instead yield a localized spatial pattern in the solution of Δ*r*_L2*/*3_(*d*) (Figure 5B_2_). Note that this sketch of our theory does not explicitly consider excitatory and inhibitory pathways, our theory can be extended to properly consider the full circuit structure (Supplementary Methods S1 and S2).

In our data, both L4 and L2/3 exhibit an absence of any spatial organization at broad (100 ∼ 200 *µm*) distances, yet a patterned, micro-clustered organization at narrow (∼ 20 *µm*) distances. We next apply our general theory for pattern formation at both of these spatial scales in our L4 – L2/3 circuit model, so as to make concrete and testable circuit predictions.

### Circuit constraints in the V1 network model for the propagation of micro-clustered structure from L4 to L2/3

To begin, in order to simplify our analysis we let the recurrent excitatory and inhibitory connectivity spatial scales obey 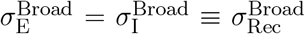. To explain the absence of any spatial patterns over broad distances, the connection widths, *σ*^Broad^, should satisfy the non-pattern forming constraint in Eq. (8):

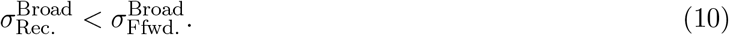

We note that since 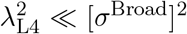 then we can neglect any L4 patterns on broad spatial scales. To test the hypothesis given in Eq. (10) we ranged our network model over 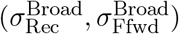 and for each pairing we computed the spatial organization of tuning preference. For a given 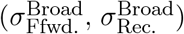 the tuning correlation curves are characterized by their peak value and spatial scale, represented by the saturation and hue of the color map, respectively (Figure 5C_1_; to range over the dense parameter space we use an associated linearized theory of our network, Supplementary Methods S2). When 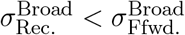 (Figure 5C_1_, above the dashed line), the network produces a spatial organization of orientation tuning with pinwheel-like neuron clusters tuned to similar preferred orientations, in disagreement with results of our L2/3 data (the red cross-mark in Figure 5C_1_ and 5C_2_). Instead, when 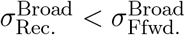 (Figure 5C_1_, below the dashed line) the network produces weak correlation over all distances and salt-and-pepper-like disorganized spatial map of preferred orientations, consistent with our L2/3 data at long distances between neuron pairs (the green check-mark in Figure 5C_1_ and 5C_2_).

To explain the micro-clustered spatial patterns over a fine spatial scale ( ∼20 *µm*), the connectivity on narrow spatial scales, *σ*^Narrow^, should place the network in the pattern-forming regime as prescribed in Eq. (9):

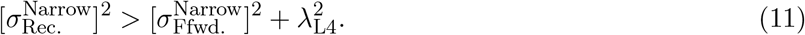

To test this hypothesis we fix (*σ*^Broad^, *κ*) (note that *κ* is the ratio of narrow component wiring in Eq. (1)) and range our model over 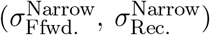. We measured the peak value and the spatial scale λ_L2*/*3_ of Gaussian fits to the distance dependent tuning curve correlations in L2/3 (Figure 5D_1_). For large 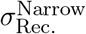 that satisfies Eq. (11), the network produces high amplitude tuning correlations over a fine spatial scale (Figure 5D_1_, above the dashed line), as well as micro-clustered patterns in the preferred orientation map, consistent with the L2/3 data measured over fine spatial scales (the green check-mark in Figure 5D_1_ and 5D_2_). In contrast, a narrow 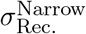 (Figure 5D_1_, below the dashed line) produces weak tuning correlations over fine spatial scale and a salt-and-pepper-like orientation map, inconsistent with our L2/3 data (the red cross-mark in Figure 5D_1_ and 5D_2_). Interestingly, for any fixed value of 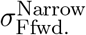, the spatial width of tuning correlations decreases with 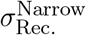, as the hue turns from red to blue in the color map (Figure 5D_1_). In fact, for sufficiently large 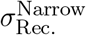 we have that 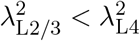, as is true for our L4 and L2/3 data. This is because lateral inhibition of recurrent connections can sharpen the spatial profile of correlations in L2/3 that are inherited from L4.

**Figure 5:**
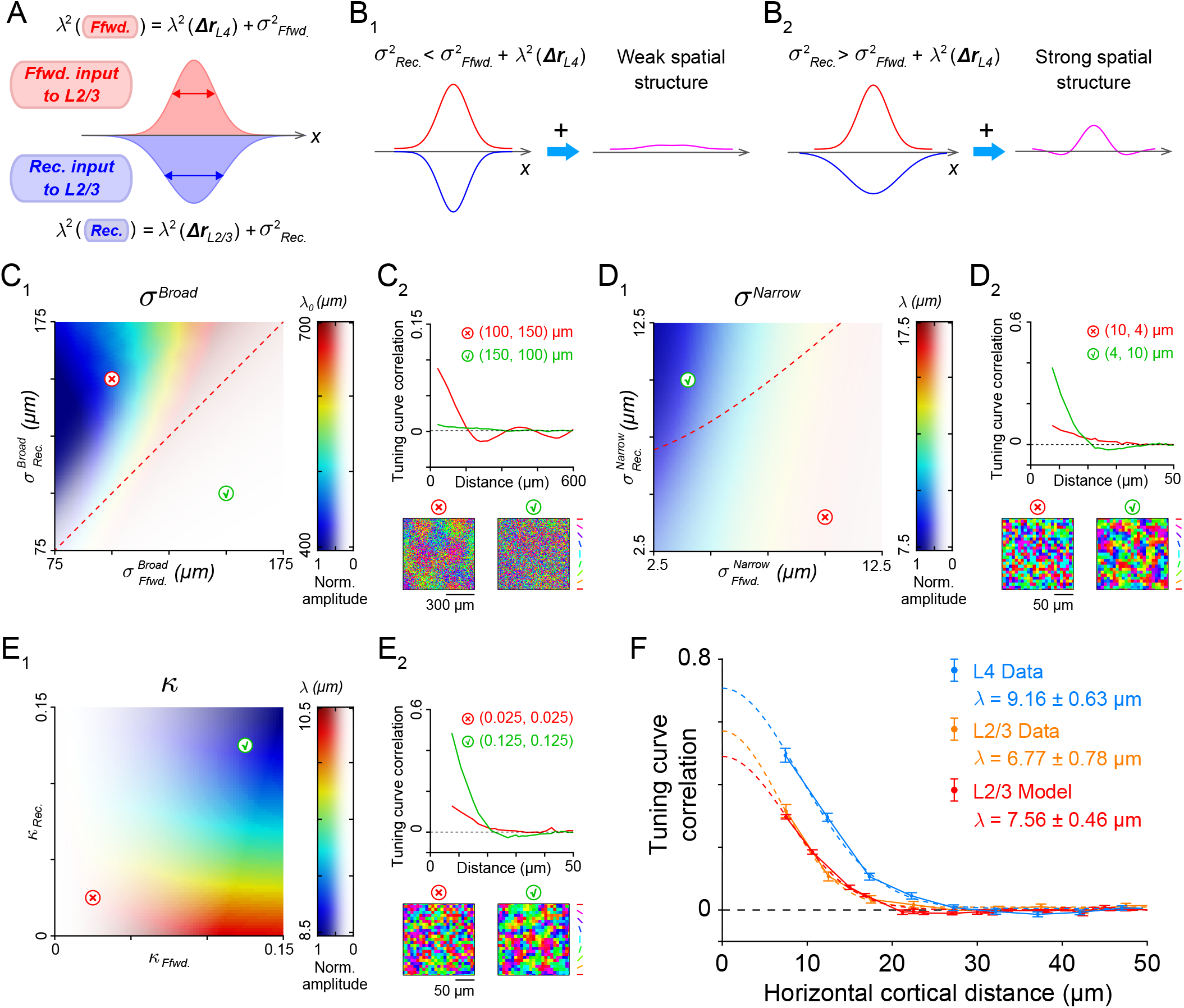
Circuit conditions for the absence/presence of spatial organization of orientation tuning over broad/narrow spatial scales in L2/3. (A) Schematic of the L2/3 neural-field model, showing Gaussian-shaped spatial profiles of excitatory feed-forward synaptic inputs (red footprint) and opposing inhibitory-dominant recurrent synaptic inputs (blue footprint) to an L2/3 neuron. The parameters *σ*_Ffwd._ and *σ*_Rec._ are the spatial widths of connectivity kernels of feedforward and recurrent projections, respectively; λ(Δ*r*_L4_) and λ(Δ*r*_L2*/*3_) are the widths of the spatial organization of L4 and L2/3 population activity, respectively. (B_1_) When the spatial widths and amplitudes of feedforward and recurrent input terms in (A) are equal, the recurrent inhibition will cancel the feedforward excitation at all spatial locations. Consequently, there will be no spatially patterned activity in L2/3. Such cancellation requires 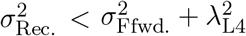. (B_2_) When recurrent connection *σ*_Rec._ becomes so broad that the condition in (B_1_) fails, the inhibitory recurrent input can not perfectly counterbalance the feedforward input. Instead, broad lateral inhibition yields a localized solution, indicating a strong, clustered spatial pattern in L2/3. (C_1_) Analysis of the network model with different combinations of 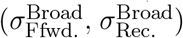. The saturation and hue of each point on the color map represents the normalized amplitude and spatial scale λ_0_ (see Supplementary Methods S1 for definition) of the distance dependent function of tuning curve correlation of L2/3 E neuron population. The dashed line marks the boundary of pattern-forming regime and non-pattern-forming regime in Eq. (10). The red cross-mark and green check-mark are two examples of 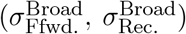 that fail/succeed to match the L2/3 data. Note that *κ* = 0 in all the simulations. We remark that all analysis for panels C_1_, D_1_, and E_1_ is done with a linear response analysis of the network activity (see Supplementary Methods S1). (C_2_) Tuning curve correlations as a function of distance (*d*) and the spatial map of preferred orientation for the examples of the red cross-mark and green check-mark in (C_1_), generalized with spiking neuron network simulation. (D and E) Same as panel (C) but with varying the connectivity on the narrow spatial scale through its width *σ*^Narrow^ (D) and amplitude *κ* (E). The dashed line in (D_1_) marks the threshold in Eq. (11). In (D),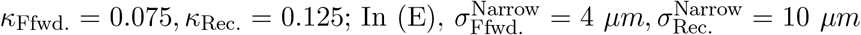 In (D) and (E), 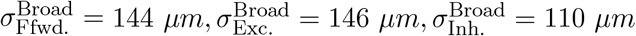. (F) The distance dependence of tuning curve correlations obtained from the network of spiking neurons models quantitatively matches the L2/3 data. The combination of parameters used are shown in Figure 6E. Error bars: data; dashed line: Gaussian fit.

Finally, we fix (*σ*^Broad^, *σ*^Narrow^) and explore how the magnitude of narrow connectivity (*κ*_Ffwd._, *κ*_Rec._) in our model network affects the spatial organization of orientation tuning (Figure 5E_1_). Small *κ*_Ffwd._ produces weak tuning correlations and a salt-and-pepper-like orientation map, regardless of *κ*_Rec._ (the red cross-mark in Figure 5E_1_ and 5E_2_). If *κ*_Ffwd._ = 0, the feedforward projection becomes a single broad Gaussian filter with a spatial scale of hundreds of microns. As a result, any fine micro-clustered organization in L4 will vanish in the feedforward input currents to L2/3, due to spatial low pass filtering from L4 → L2/3. By contrast, when *κ*_Ffwd._ is sufficiently large, the spatial width of tuning correlations (represented by hue of the color map) decreases with *κ*_Rec._ (the green check-mark in Figure 5E_1_ and 5E_2_), consistent with our previous finding that recurrent wiring will sharpen inherited spatial correlation structure. In total, a non-zero *κ* is necessary for the model to produce a high level of tuning correlation in L2/3 over a fine spatial scale.

The guidance of these theoretical predictions allowed us to fine tune the *σ*^Broad^, *σ*^Narrow^ and *κ* of the feedforward and recurrent E and I connections in our spiking network model (see Figure 6C and Methods for model parameter values) so as to quantitatively match the distance dependent function of tuning curve correlation in the L2/3 data (Figure 5F). These model circuitry parameters provide clear, testable predictions for synaptic wiring in L4 and L2/3 of mouse V1. The remaining sections of our study are devoted to testing these predictions through several, complementary experiments.

### Circuit mapping verified the model predictions of intracortical connectivity

To measure the spatial scales of feedforward and recurrent connectivity in L2/3 we analyzed a previously collected dataset of rabies-traced presynaptic ensembles of individual L2/3 pyramidal (E) neurons in mouse V1 (Rossi et al., 2020) (Figure 6A). We sorted the presynaptic neurons into excitatory populations from L4 (i.e. feedforward excitation), excitatory populations from L2/3 (recurrent excitation) and inhibitory populations from L2/3 (recurrent inhibition). We collected the coordinates (Δ*x*, Δ*y*) of recorded presynaptic neurons relative to the location of their postsynaptic L2/3 E neurons (n = 17, Figure 6B). Finally, we used maximum likelihood estimation to fit the parameters *σ*^Broad^, *σ*^Narrow^ and *κ* of the sum-of-Gaussians probability distribution (Eq. 1) for the three types of connections (results summarized in Figure 6C, upper panel).

We first compared the feedforward and recurrent connections on broad spatial scales, and found that 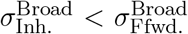 (110.6 *µm* < 143.6 *µm*). These estimates are consistent with our theoretical predictions (Eq. 11) of the connectivity needed to avoid any long-range spatial organization of tuning preference in L2/3. Indeed, to match the tuning correlation curves estimated from population data (Figure 5F), our network model required *σ*^Broad^ estimates that closely match the anatomical data (compare the values in Figure 6C, and overlapping curves of data and model at long distance in Figure 6D). In addition, the rabies tracing dataset also validated the ‘like-to-like’ feature (Δ preferred orientation) dependent wiring rule (Figure S6), and the statistical independence of the spatial-dependent term and the feature-dependent term in the model (Figure S7), as required by Eq (3).

We next considered connectivity over narrow spatial scales, and estimated that 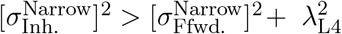 (Figure 6C, upper panel). This is consistent with the theoretical condition of pattern formation over fine spatial scales (Eq. 11). We note that the 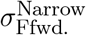 estimated from the anatomical data is broader than that in the model (Figure 6C, compare upper and lower panel, 12.20 *µm* v.s. 5.34 *µm*). However, the largest disagreement between the model and anatomical data is that the values of the estimated *κ*s from data are an order of magnitude weaker than that required for our model to match the tuning curve correlations (compare the *κ* values in Figure 6C, and data and model curves at short distances in Figure 6D).

One possible reason for such a mismatch between theory and experiment is that monosynaptic rabies tracing only reports the presence of connections between pre- and post-synaptic neurons, but it cannot inform about the strengths of these connections. Connections between nearby neurons (or vertically aligned postsynaptic neurons for feedforward projection) may have more synaptic contacts and hence stronger effective post-synaptic potentials than that of distant neuron pairs (Cossell et al., 2015; Li et al., 2012; Yu et al., 2009). Alternatively, ephaptic coupling would increase the effective post-synaptic potential for connections from very nearby presynaptic neurons (Anastassiou et al., 2011). Regardless, the functional neuronal interactions may show a sizable narrow component, while the anatomical wiring only shows a weak bias to narrow connectivity.

**Figure 6:**
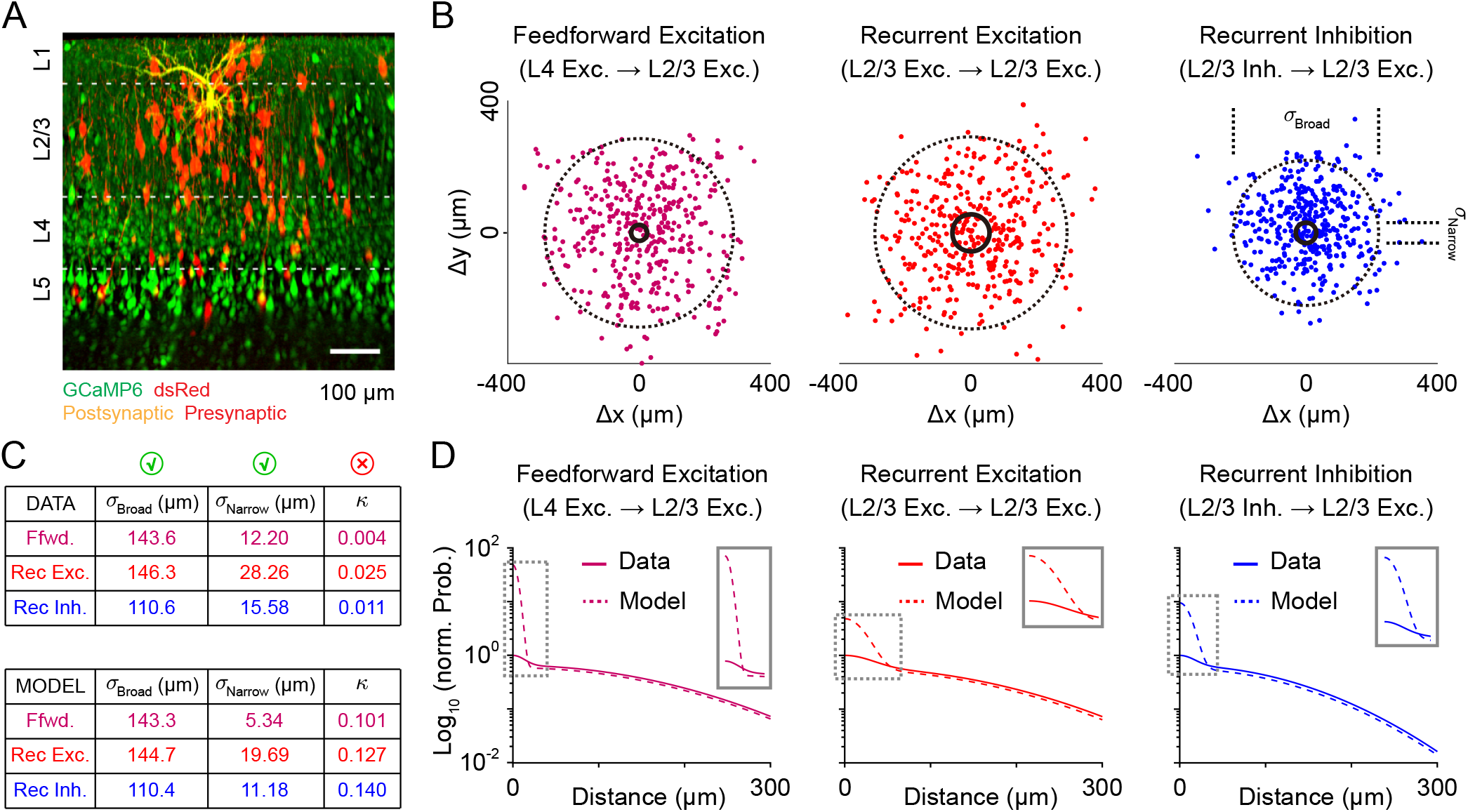
Anatomical mapping of the spatial profiles of excitatory and inhibitory connections to L2/3 pyramidal (E) neurons in mouse V1. (A) The presynaptic population of an example postsynaptic L2/3 pyramidal neuron. Montage of *in vivo* two-photon sagittal projections showing the postsynaptic neuron: yellow; the presynaptic neurons across multiple layers: red, expressing dsRed; other pyramidal neurons: green, expressing GCaMP6. Modified from (Rossi et al., 2020). (B) Relative coordinates (Δ*x*, Δ*y*) of presynaptic populations of feedforward excitation (magenta, n = 410), recurrent excitation (red, n = 418) and recurrent inhibition (blue, n = 406), stacked vertically (over z). The origin represents the location of postsynaptic L2/3 pyramidal neurons (n = 17, overlapped). Circles: contours of the estimates of *σ*^Broad^ and *σ*^Narrow^. (C) Top: *σ*^Broad^, *σ*^Narrow^ and *κ* of the sum-of-Gaussians probability distribution function for three types of connections, estimated by maximum likelihood estimation (MLE) from the samples in (B). Bottom: the parameters used for the model in Figure 5F. (D) Comparison of the spatial profile of the three types of connections between data estimation (solid) and model (dashed) in Figure 5F. Inset: zoom in of the mismatch on the narrow spatial scale.

**Figure 7:**
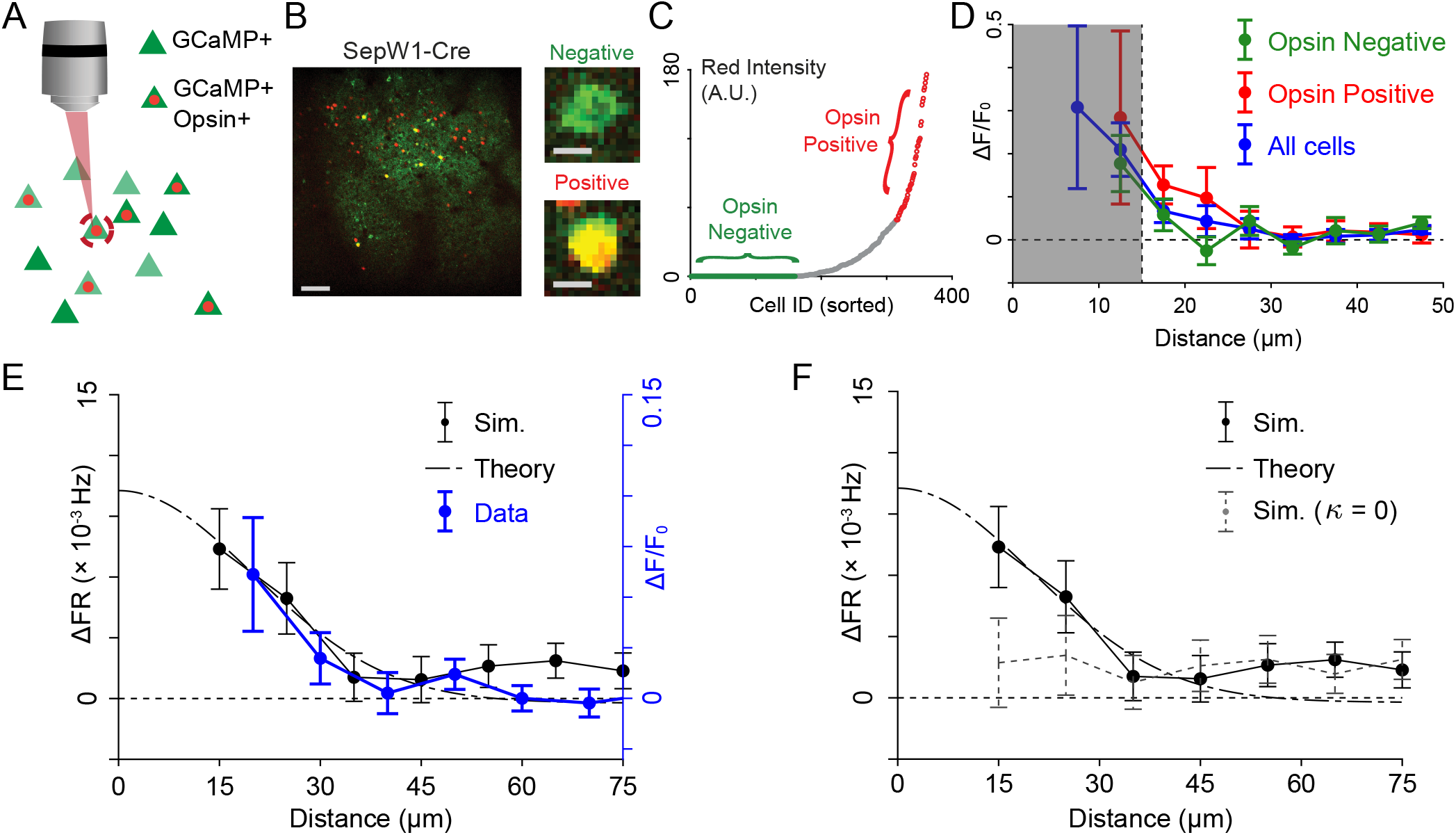
Functional mapping the spatial profile of L2/3 E → L2/3 E excitatory recurrent connections in mouse V1 via single cell multi-photon optogenetics. (A) Schematic of photo-stimulation with single cell resolution. (B) Sparse expressing light sensitive opsin; neurons with postive expression are in yellow, and neurons with weak opsin expression are in green. Scale bar: zoomed out, 100 *µm*; zoomed in, 10 *µm*. (C) Opsin-negative neurons and opsin-positive neurons sorted in the ensemble of L2/3 pyramidal neurons. (D) The spatial profile of trial-averaged Δ*F*/*F*_0_ change of L2/3 pyramidal neurons relative to the location of photo-stimulated cell, for ensembles of opsin negative (green), positive (red) and all cells (blue). Responses closer than 15 *µm* are excluded from further analysis (gray zone). Average of n = 112 single cell stimulation, for 4 field of views (FOVs) in 4 mice. (E) Comparison of the spatial profile of trial-averaged firing rate change (ΔFR(d)) in the spiking network model (average of 15000 trials), and in the linear response theory (see Supplementary Note) (both using the same model parameters as in Figure 5F and Figure 6C), and the trial-averaged Δ*F*/*F*_0_ change of all cells in the data (same as in (D)). (F) Comparison of ΔFR(d) for the results from spiking network model, linear response theory, and the alternative spiking network model that removed the narrow component of all intracortical connections (i.e. *κ* = 0).

To probe the spatial wiring of functional synaptic connectivity we employed *in vivo* single cell resolution multi-photon optogenetics in L2/3 of mouse V1 (Oldenburg et al., 2024). The temporally focused optical system and the optimized fast opsin (‘ChroME’) we developed (Mardinly et al., 2018) enable optogenetic activation of individual neurons with cellular resolution at extremely high temporal resolution (Figure 7A). However, a well known caveat of single cell resolution optogenetics is that the focus of the laser beam could activate unintended neurons that are near the target neuron. This is of particular concern for our study, since the range of the point spread function of the laser beam is comparable to the scale of *σ*^Narrow^ (Mardinly et al., 2018), and could seriously impact the measurement of Δ*F*/*F*_0_ for neurons close to the photo-stimulated cell. To minimize any off-target effects we used a sparse expressing opsin in SepW1-Cre transgenic mice to curtail the co-expression of opsin in the surrounding L2/3 population (Figure 7B, 7C). Indeed, photo-activation of opsin-negative neurons was minimal compared to that of opsin-positive neurons.

We performed simultaneous photo-stimulation and two-photon calcium imaging of L2/3 pyramidal neurons, and we targeted only a single opsin-positive neuron for stimulation. This was done so that the cortical response would be relatively weak, and we may assume that it reflects only mono-synaptic connections from the stimulated to recorded neurons, unlike the case when multiple neurons are targeted (Marshel et al., 2019; Oldenburg et al., 2024). For each trial, the target photo-stimulated cell evoked 10 spikes within 1 second, and the trial-averaged Δ*F*(*t*)/*F*_0_ of surrounding L2/3 pyramidal neurons were recorded. We measured the distances between the recorded neurons and the target neuron, and constructed the average spatial profile of the functional connectivity of L2/3 E → L2/3 E excitatory recurrent connections. There was little influence on recorded neurons that were > 30 *µm* from the stimulated neuron, while there was significant activation of neurons 15 ∼ 25 *µm* from the stimulated neuron (Figure 7D, all curves). And satisfyingly, the spatial profile of Δ*F*/*F*_0_ activation for ensembles of opsin-negative cells only, opsin-positive cells only, as well as all cells did not show any significant differences (Figure 7D, green, red and blue curves, respectively). To be conservative, we excluded any cells that were closer than 15 *µm* from the stimulated target in further analysis (Figure 7D, gray zone).

To validate the connectivity parameters *σ*^Narrow^ and *κ* we hypothesized in our model, we mimicked the single cell optogenetic activation in our spiking network model by evoking spikes in L2/3 E neurons with the same protocol as in the experiments. The model used the same connectivity parameters which provided a quantitative match to the spatial scales of orientation tuning (Figure 5F and Figure 6C). The spatial profile of the relative trial-averaged firing rate change (ΔFR(d)) is also a near quantitative match to the results of the photo-stimulation data (Figure 7E, compare the black and blue error bar curves). Apart from the simulations of the network of spiking neuron models, the ΔFR(d) computed from closed-form mathematical theory of linear response neural field (see Supplementary Note) also agree with results of simulation and data (Figure 7E, dashed curve). These agreements provide a validation of the *σ*^Narrow^ and *κ* of L2/3 E → L2/3 excitatory recurrent connection used in the model. As a comparison, we performed the same activation in the spiking network model, but removed the narrow component of all intracortical connections (*κ* = 0). In this case ΔFR(d) has no peaked response over distances < 20 *µm* (Figure 7F, dashed error bar), which further points to the necessity of a narrow spatial component of recurrent wiring in L2/3.

**Figure 8:**
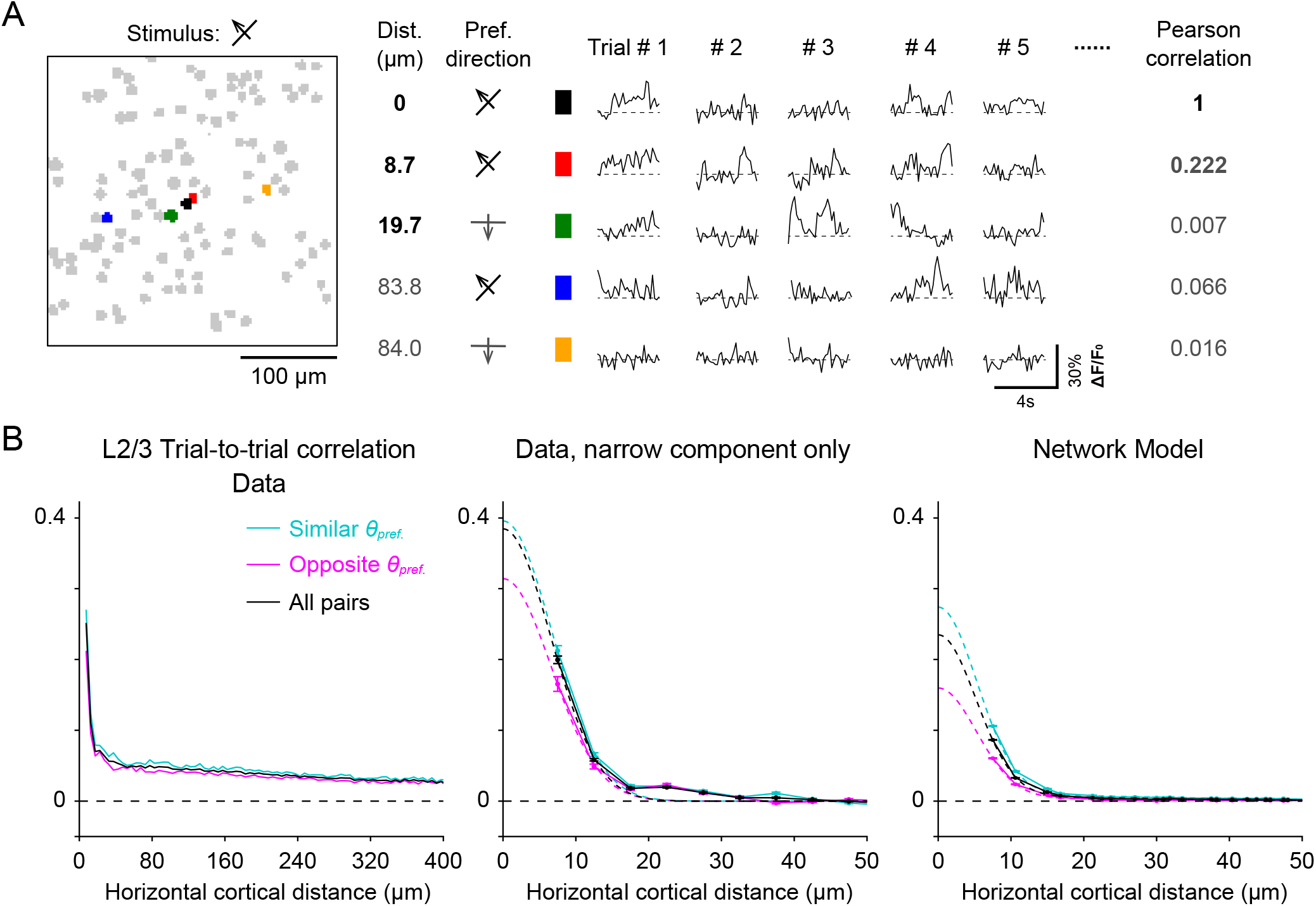
The spatial organization of noise correlations in L2/3 of mouse V1. (A) The Δ*F*/*F*_0_(*t*) traces of five example ROIs in response to drifting grating stimuli in one specific direction (135^*o*^) over multiple trials. The black ROI in the center of the field of view is set as the reference neuron. For the other 4 example ROIs labeled in different colors, the table reports their horizontal cortical distance to the reference, preferred direction, response over 5 example trials, and the Pearson correlation of responses over all 250 trials to the reference ROI (i.e. trial-to-trial correlation). Only the red ROI with similar preferred feature and close distance to the reference neuron has strong noise correlations with the reference neuron. (B) The average trial-to-trial noise correlation function over horizontal cortical distance in L2/3 data and model. The blue, pink and black curves show noise correlations computed from pairs of L2/3 neurons with similar preferred directions, opposite preferred directions, and all pairs, respectively. Error bars: data; dashed line: Gaussian fit. Left: the noise correlation data over narrow and broad spatial scales. Middle: Data after removal of the spatially broad, conjectured latent shared source of external correlations. Right: the spatial organization of noise correlations function in the spiking network model (using the same parameter as in Figure 5F, 6C and 7E-7F), which qualitatively matches the narrow component of the L2/3 data.

### The narrow spatial profile of L2/3 noise correlations supports circuit predictions

In the previous sections we characterized the trial-averaged neuronal responses driven by visual stimuli, and used the spatial organization of orientation tuning to constrain models of L4 – L2/3 circuit in mouse V1. Another well established expression of circuit architecture is the trial-to-trial covariability of neuronal response, often termed noise correlations (Doiron et al., 2016; Hennequin et al., 2018; Ocker et al., 2017; Renart et al., 2010; Rosenbaum et al., 2017; Shadlen and Newsome, 1998; Urai et al., 2022). Briefly, if a pair of neuronal responses show shared trial-to-trial variability it suggests that the neurons share significant pre-synaptic inputs, and it is the trial-to-trial fluctuations of these inputs that correlates post-synaptic responses. Evidence for this relationship between circuitry, tuning, and variability already exists in L2/3 of mouse V1, as pairs of E neurons that are connected show both higher correlations in their tuning preferences as well as high noise correlations (Cossell et al., 2015; Ko et al., 2013). The narrow circuitry that supports the fine scale spatial organization of trial averaged tuning response should produce a fine scale organization of noise correlations (Rosenbaum et al., 2017). We test this prediction in this final section.

To properly estimate noise correlations over the L2/3 population we measured the calcium traces Δ*F*(*t*)/*F*_0_ of identified neurons in awake mice, in response to sinusoidal drifting grating stimuli over 250 trials in one of four motion directions. It was necessary to reduce the number of stimulus conditions to properly sample the within condition variability of population response. An example pair of neurons with similar preferred directions and a short cortical distance showed high noise correlations (example in Figure 8A, compare the black and red ROIs), while example pairs with mismatched preferred directions or longer distances lead to lower correlations (example in Figure 8A, other three ROIs). To investigate the population as a whole we computed the average noise correlations conditioned on the horizontal cortical distance between L2/3 neuron pairs (Figure 8B, black curve).

The spatial profile of noise correlations contains one fast decaying narrow component, and another slowly decaying broad component (left panel, Figure 8B). We conjecture that the broad component might represent shared sources of latent variability that are external to the L4 – L2/3 network (such as inputs related to overall brain state), since low-dimensional sources of variability would increase correlations at all distances (Musall et al., 2019; Stringer et al., 2019). It is unlikely that the narrow component of noise correlations is due to brain state fluctuations, as they would not have any fine spatial specificity. To isolate the narrow spatial component we fit the noise correlations against distance curves with the sum of one narrow and one broad Gaussian-shaped functions, and subtracted the broad component of the fit. The residual, narrow component exhibited strong noise correlations over a fine spatial scale, which holds for overall neuron population, as well as subpopulations with similar or opposite orientation preferences (middle panel, black, cyan and magenta curves respectively, Figure 8B). This is similar to the structure of tuning curve correlations (Figures 1, 3 and 5). In total, our analysis of population-wide trial-to-trial variability provides further evidence for the spatially narrow circuitry component in L4 – L2/3 V1 circuit.

## Discussion

Our *in vivo* calcium imaging and population analysis examined awake mouse V1 over a fine spatial scale that is often overlooked. Our analysis established a ‘micro-clustered’ organization of orientation preference in both L4 and L2/3 spanning ∼ 20 *µm*, as well as confirmed the absence of any spatial organization over longer distances (Figures 1 and 2). These results are consistent with past studies of functional micro-architectures in mouse V1 (Kondo et al., 2016), but at odds with the traditional notion of a true “salt-and-pepper” organization in rodent V1 (Bonin et al., 2011; Han et al., 2019; Kaschube, 2014; Ohki et al., 2005; Ohki and Reid, 2007), or broad spatial clustering in L2/3 spanning over hundreds of microns (Ringach et al., 2016). The combination of our ground-truth measurement by neuropil-free nuclear-targeted GCaMP, as well as the cytosolic GCaMP datasets with optimal neuropil subtraction, provide substantial evidence for the ‘micro-clustered’ organization in L2/3 (Figure 3). Our results underscore the role of nuclear-targeted GCaMP as a faithful validation of measurements using other methods (Barchini et al., 2018), as well as the importance of optimal neuropil correction in two-photon calcium imaging with cytosolic GCaMP. Indeed, an inappropriate neuropil correction – with insufficient or excessive subtraction factor *α* – would produce an artifactual organization of long-range tuning similarity across tens of microns (see cases of *α* < 1 and *α* > 1 in Figure 3H). In cases where nuclear GCaMP measurement is not viable – for example, due to difficulty in expression in target neurons (e.g. in deeper L4 neurons), or observing timescale sensitive neuronal activities – our work stresses the importance of checking the credibility of results with different subtraction factors *α* for cytosolic GCaMP datasets.

The central hypothesis about V1 circuitry in our network model is a narrow component of synaptic wiring that has a comparable spatial scale to the micro-clustered organization of orientation preference. One possible developmental origin of this narrow scale for circuitry is ontogenetic columns formed during corticogenesis (Singer et al., 2019). According to the radial unit hypothesis, during embryonic development, neurons from the same neuronal progenitor cell will migrate and form aligned strings spanning cortical layers, termed ontogenetic columns (Rakic, 1988). Neurons within the same ontogenetic column (termed ‘sibling’ or ‘sister’ neurons) in mouse neocortex tend to preferentially establish chemical synapses with each other, rather than other neighboring non-sibling neurons. This process is possibly mediated by electrical coupling (gap junctions) formed transiently during very early stages of development (Yu et al., 2009). These interconnected sibling neurons tend to show similar feature selectivity, including orientation preference of visual stimuli in mouse V1 (Li et al., 2012). However, it still remains unclear how the strength (i.e. *κ*) and the spatial spread (i.e. *σ*^Narrow^) of these narrow scale connections among sibling neurons are regulated. Future work should assess how the neuronal developmental process, as well as synaptic plasticity mechanisms before eyes opening, could shape narrow scale circuitry in V1 altogether.

A counter-intuitive consequence of our micro-cluster theory is that a very small number of physically close neurons to a post-synaptic target will nevertheless have a large influence on post-synaptic response. This outsized micro-circuit influence must persist in the face of the rest of the global macro-circuit which contains the vast majority of pre-synaptic activity. We argue that this micro-circuit influence is due to the distinct regimes of E - I balance over different spatial scales (sketched in Figure 9).

Let the total number of pre-synaptic inputs to a neuron be *K*. We take *K*_micro_ << *K* of these inputs to be from the surrounding micro-circuit and the remaining *K* − *K*_micro_ inputs to be from the macro-circuit. The challenge here is for the input currents onto a post-synaptic neuron from both the micro- and macro-circuits to be of comparable and moderate magnitude (by moderate we mean *𝒪*(1) even for large *K*). In a balanced network framework it is typical to let synaptic strength be large (i.e. 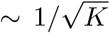 as opposed to 1/*K*), so that for a large pre-synaptic pool *K* the total input could, in principle, diverge (*K* inputs each with strength 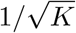 could give 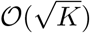 total input and not *𝒪*(1) as needed). Following classic work in the theory of balanced networks (Shadlen and Newsome, 1998; van Vreeswijk and Sompolinsky, 1998) we assume that for the macro-circuit we have a balanced cancellation between feedforward and recurrent inputs, so that even for large *K* the total input remains of moderate magnitude (Figure 9, green box). In a spatially distributed network, such a balanced cancellation must be operative at all distances so that no spatial organization can emerge over the macro-scale (Rosenbaum et al., 2017). By contrast, over the micro-scale if we take 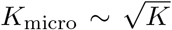 then synaptic strength is still very large, yet the small size of the micro-circuit allows the total input to be moderate without a need for balancing (Figure 9, orange box). As a result, a spatial imbalancing will create patterned organization of activity over micro-scale distances (Rosenbaum et al., 2017).

**Figure 9:**
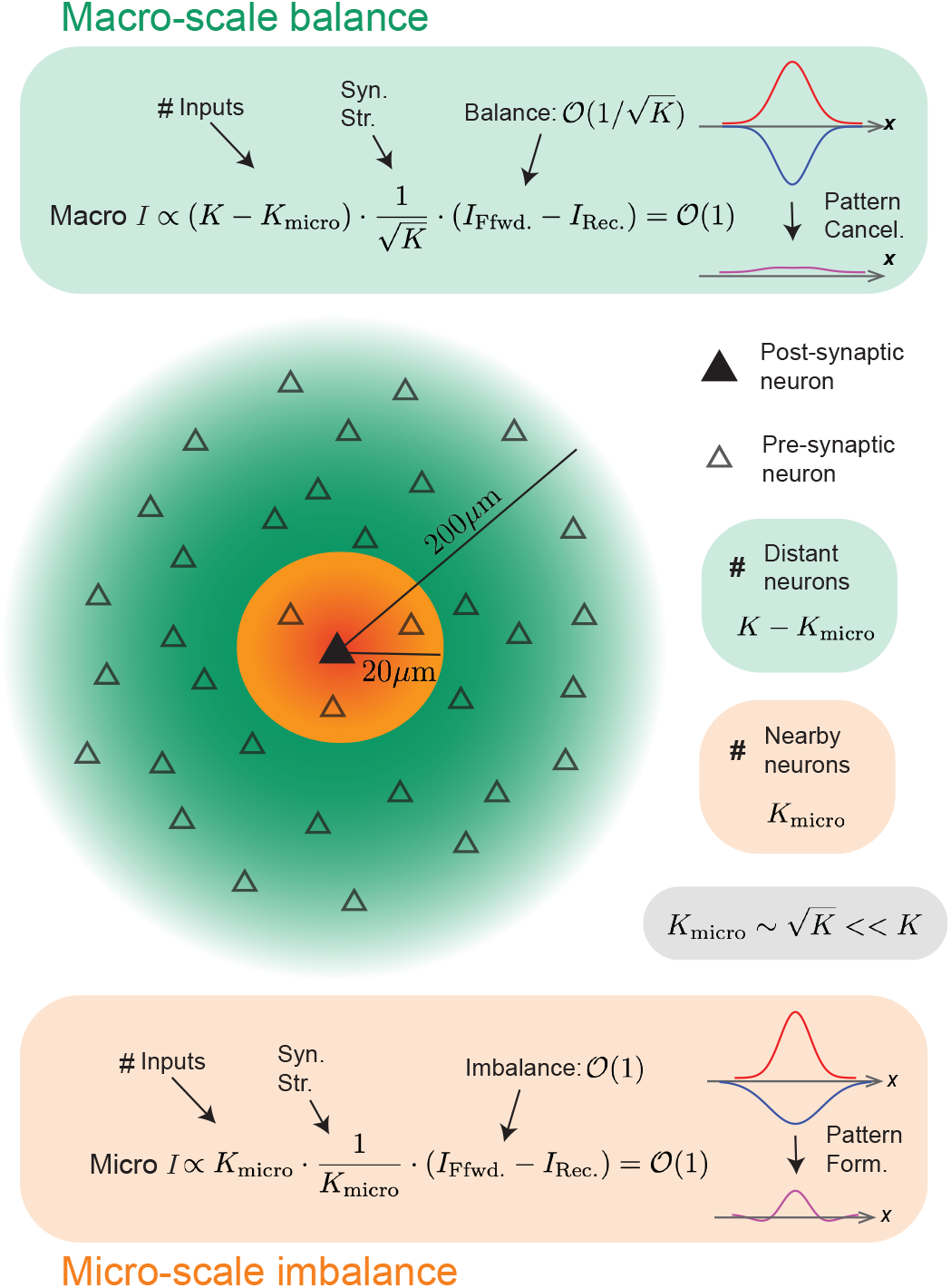
Illustration of macro-scale balance and micro-scale imbalance in L2/3 of mouse V1. Consider the mean current given to a post-synaptic neuron (filled triangle) from the macro-population located 20 *µm* – 200 *µm* from the post-synaptic neuron (open triangles in green region) and the micro-population located within 20 *µm* of the post-synaptic neuron (open triangles in orange region). Take the total number of inputs to the post-synaptic neuron to be *K*, with *K*_micro_ << *K* being within the micro-population, and the remaining *K* − *K*_micro_ being in the macro-population. The colored boxes decompose the mean current into the product of: 1) the number of pre-synaptic inputs, 2) the synaptic strength, and 3) the difference between feedforward and recurrent inputs. Over the macro-scale, in order for the product of these quantities to be well behaved (i.e be 𝒪(1)) and have large synaptic strengths relative to population size (i.e synapses scale as 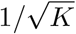, we require that the feedforward and recurrent inputs balance one another (see green box). This results in no pattern formation over macro-scales. Over the micro-scale we take the synaptic strength to scale as 1/*K*_micro_, where we take 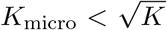 (implying larger synaptic strength). This gives a well behaved mean input even when the feedforward and recurrent inputs are in imbalance with one another, yielding pattern formation over the micro-scale (see orange box).

In combining these two results together we see that it is the balanced cancellation over the macro-scale that yields moderate inputs (of *𝒪*(1) magnitude) and this permits the imbalanced micro-scale inputs to be comparable in influence (of *𝒪*(1) magnitude) despite *K*_micro_ << *K*. However, if there was an imbalance over the macro-scale, then it would dominate all activity and there would consequently be no possibility of a patterned organization over the micro-scale. In other words, it is the balanced activity over the macro-scale that allows imbalanced activity over micro-scales to compete with the much larger macro-synaptic field. Similar arguments of networks having a global balance with selective local imbalance have been made for networks with assembly structure (Litwin-Kumar and Doiron, 2012; Roudi and Latham, 2007; van Vreeswijk and Sompolinsky, 2005).

Our circuit model explains the narrow spatial organization of orientation preference by assuming a narrow spatial scale of feedforward and recurrent coupling (*κ* > 0). In essence we have explained functional organization through an assumed circuit structure. An alternative approach would be to assume a biological rule, that is yet to be discovered, which organizes orientation preference to be clustered over fine spatial scales. This spatial organization would then not require a narrow scale of synaptic coupling. Rather, the combination of spatial clustering of orientation preference and a like-to-like feature-based wiring would now predict a narrow spatial scale coupling, as suggested in our experimental data (Figures 6 and 7). Thus, in this alternative model we would have explained a circuit structure through an assumed functional organization. This ‘chicken-or-egg’ dilemma of the causal connections between structure and function in network neuroscience is a recurring theme (Bassett et al., 2018). One possible way to distinguish between the two models would be to consider the functional organization (or lack thereof) of noise correlations over narrow spatial scales (Figure 8B). The alternative funtion to circuit model would require sufficiently narrow excess spatial coupling, due only to like-to-like feature-based wiring and orientation clustering, so as to create narrow spatial scale noise correlations (Figure 7E). However, it would also require sufficiently broad feature wiring to allow nearby neurons with opposite feature preference to still show narrow noise correlations (Figure 8B, magenta curve). This may require a fine tuning of the synaptic wiring rules.

Finally, we consider possible functional consequences of micro-clustered organization in mouse V1. The orientation preference maps in V1 from higher mammals are tiled with pinwheel structures, where neurons with different preferred orientations are circularly arranged around a central focus (Ohki et al., 2006). Every spatial location in the visual field is then covered by the receptive fields of neurons in one corresponding pinwheel unit, so that all possible preferred orientations have some representation. However, visual stimuli that are smaller than the size of the pinwheel unit may not have coverage for certain orientations, and consequently will have degraded coding. Thus, the size of a single pinwheel unit structure (or the density of pinwheel tiling) in the orientation map of V1 set the limits of visual acuity (Srinivasan et al., 2015).

To be more specific, the Nyquist spatial frequency of periodically tiled clusters of neurons with similar preferred orientation, *f*_Nyq._, should be greater than the visual acuity of the animal (*f*_max._, the maximum spatial frequency of distinguishable inputs):

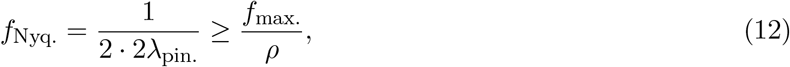

where λ_pin._ is the spatial scale of single neighboring orientation cluster, so 2λ_pin._ is the wavelength of orientation map; and *ρ* is the magnification factor from the visual field to V1.

In mouse, previous studies have shown that *f*_max._ ∼ 0.56 (cyc/deg.) and *ρ* ∼ 20 (*µm*/deg.) (Van Hooser et al., 2005). This sets λ_pin._ ≤ ∼ 9 (*µm*/cyc.), which agrees with the scale of micro-clustered organization that we report. That is to say, the micro-clustered organization of orientation preference in rodents is possibly a severely degraded version of the pinwheel structure seen in V1 of higher mammals. This is a joint consequence of the small area of V1 (∼1 mm^2^) and a low magnification factor of the visual field in mice, as well as a low visual acuity (*f*_max._) of mice due to the large size of their receptive fields. In mouse V1 neighboring pairs or triplets of neurons must encode the same region of the visual field, as the neuron population in a full pinwheel structure in the V1 of higher mammals. Consequently, the orientation encoding accuracy of neurons in micro-clusters in mouse V1 will certainly be much lower than that of a full pinwheel. However, it will be higher than that of one single neuron in a strict ‘salt-and-pepper’ organized orientation map.

In sum, our study has identified a fine spatial scale (∼ 20 *µm*) micro-cluster organization of orientation selectivity in mouse V1. Our modelling efforts predicted the specific circuit structure at both broad and narrow scales required to account for this finding. While we have speculated on the functional impact of such a micro-structure, there remain many open questions about how small groups of nearby neurons can support neuronal processing that is typically associated with large populations of neurons that span whole brain regions.

## Methods

### Mice

All animal procedures and protocols were approved by the Institutional Animal Care and Use Committee at the University of California, Berkeley (AUP-2020-06-13343, AUP-2014-10-6832-2). Wild-type mice (C57BL/6J) and Scnn1a-Tg3-Cre mice older than 2 months were used for in vivo functional imaging of L2/3 and L4 neurons in V1, respectively. Male and female mice were equally represented, and no sex differences were observed.

### Sample preparation: virus injection and cranial window implant

Craniotomy and virus injection were performed on adult mice following established procedures (Sun et al., 2016). Buprenorphine (0.33 *µ*L/g) was administered subcutaneously at the start of the surgery. Mice were anesthetized with 1%-2% isoflurane in oxygen throughout the surgery.

A 3-mm craniotomy was made over the primary visual cortex (V1) for optical access. The dura was left intact. Virus injection was performed using a glass pipetted beveled at 45^*o*^ with a 15-20 *µm* opening and backfilled with mineral oil. Viral solution was loaded and injected into the cortex 300-350 *µm* below dura with a hydraulic manipulator (MO10; Narishige). 30 nL of AAV2/1-syn-GCaMP6s (∼3×1013 GC/mL) or AAV2/1-Syn-H2B-GCaMP6s (∼5×1012 GC/mL) solution were injected into wild-type mice to express GCaMP6s in the cytosol or the nucleus, respectively. 30 nL of AAV2/1-syn-FLEX-GCaMP6s (∼1×1013 GC/mL) solution was injected into V1 of Scnn-Tg3-Cre mice to express GCaMP6s cytosolically in L4 excitatory neurons in V1. A cranial window made of a No. 1.5 coverslip was placed over the craniotomy and glued in place with Vetbond (Vetbond; 3M). A titanium headbar was attached to the skull with cyanoacrylate glue and dental acrylic. Meloxicam was given (1 *µ*L/g) subcutaneously after the surgery.

### *In vivo* two-photon imaging

In vivo two-photon imaging was performed 2-4 weeks after the surgery. All imaging recordings were carried out on awake mice that were habituated for head fixation and body constraint. Two-photon calcium imaging was conducted using a Thorlabs Bergamo^®^ II multiphoton microscope. Hardware control and data acquisition were performed using the ThorImage software. Fluorescence from GCaMP6s was excited by a titanium–sapphire laser (Chameleon Ultra II, Coherent Inc.) at 920 nm. The laser beam was focused by a Nikon 16×, 0.8 NA water-dipping objective. The emitted two-photon fluorescence was collected by the same objective, spectrally filtered by a bandpass filter (525/50 nm) and detected by a photomultiplier tube (PMT).

The post-objective power was varied between 20 and 80 mW, depending on the imaging depth. Typical images had 1024 × 1024 pixels at 0.7-1 *µm*/pixel and a frame rate of 7.6 Hz. Data of L2/3 neurons were acquired from 34 imaging planes of 10 mice at imaging depths between 110 and 290 *µm* below dura. Data of L4 neurons were acquired from 44 imaging planes of 5 mice at imaging depths between 220 and 460 *µm* below dura.

### Visual stimulation

Visual stimulus was generated by custom-written MATLAB^®^ codes using *Psychtoolbox* (Brainard and Vision, 1997) and displayed on a screen (Samsung Galaxy Tablet). To measure the orientation tuning properties of neurons, circular sinusoidal drifting gratings were presented on a screen 8 cm away from the right eye of the mouse, covering 70^°^ × 70^°^ of visual space. The gratings was displayed in the blue channel of the screen to stimulate the blue opsin of the mouse retina (Z. Tan et al., 2015). The gratings had 100% contrast, a spatial frequency of 0.07 cycles per degree, and a temporal frequency of 2 cycles per second. The onset of each visual stimulus was marked by the appearance of a small patch of bright pixels that was detected by a photodetector (Thorlabs, PDA10A2) to trigger image acquisition.

For micro-clustering measurements, drifting gratings were presented in 12 directions (at 30^*o*^ increments). Each direction was repeated 10 times in a pseudorandom sequence. Each stimulus lasted for 10 seconds, with a uniform blue screen displayed for the first 3 and last 2 seconds and a drifting grating presented for the middle 5 seconds.

For trial-to-trial noise correlation measurements, gratings were presented in 4 directions ({45, 135, 180, 270} deg.). Each direction was repeated 250 times in a pseudorandom sequence. Each stimulus lasted for 6 seconds, with a uniform blue screen displayed for the first 3 seconds and a drifting grating presented for the last 3 seconds.

### Image processing, signal extraction and tuning curve analysis

Calcium imaging data was processed by custom-written MATLAB^®^ codes. Time-lapse image sequences were registered with a rigid-motion correction algorithm (Pnevmatikakis and Giovannucci, 2017). Regions of interest (ROIs) representing individual cortical neurons were detected and outlined for each image sequence by *Suite2p* (Pachitariu et al., 2017). Overlapped ROIs were manually removed in *Suite2p* GUI before further processing. Raw fluorescence signal for each neuron, *F*(*t*)_raw_, was extracted as the average value of the pixels within the ROI representing the neuron. To extract the neuropil signal, a square neuropil mask extending 30 *µm* from each neuron’s center (i.e., the center of mass of all pixels within the ROI) was defined. The neuropil signal of each neuron, *F*(*t*)_neuropil_, was the average value of pixels within the neuropil but outside the ROIs identified as neurons. Defining the baseline fluorescence of the neuropil, *F*_0,neuropil_, as the mode of signal distribution of *F*(*t*)_neuropil_, we then calculated the neuropil contamination signal as Δ*F*(*t*)_neuropil_ = *F*(*t*)_neuropil_ − *F*_0,neuropil_. We subtracted the neuropil contamination signal, weighted by a neuropil coefficient *α*, from the raw fluorescence signal of each neuron to acquire the neuronal calcium signal as *F*(*t*) = *F*(*t*)_raw_ − *α* · Δ*F*(*t*)_neuropil_. Defining the baseline fluorescence of the neuron, *F*_0_, as the mode of signal distribution of *F*(*t*), we calculated the calcium transient as Δ*F*(*t*)/*F*_0_ = (*F*(*t*) − *F*_0_)/*F*_0_. For simplification, *α* is uniform for all cells.

For micro-clustering measurements, the responses of each ROI to each stimulus direction were defined as the average Δ*F*/*F*_0_ during the time window of stimuli presentation across totally 10 trials. Thus, we defined the direction tuning curve *r*(*θ*) across totally 12 stimulus directions for each ROI. For ROIs considered to be responsive (maximum Δ*F*/*F*_0_ of tuning curve above 0.1), and with significantly different responses across stimuli directions (one-way ANOVA test, *P* < 0.05), we fit their direction tuning curves *r*(*θ*) with a bimodal Gaussian function:

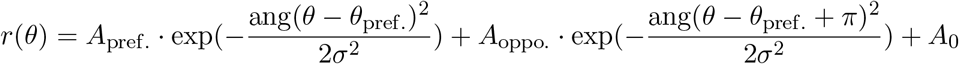

where *θ*_pref._ is the preferred direction, *A*_pref._ and *A*_oppo._ are the responses at the preferred direction and the opposite preferred direction (i.e. *θ*_pref._ − *π*) respectively. The function ang(Δ*θ*) = min(|Δ*θ*|, |Δ*θ* − 2*π*|, |Δ*θ* + 2*π*|) wraps periodic angular values onto the interval [0, 2*π*]. *A*_0_ is the constant offset. *σ* is the width of each Gaussian peak. The preferred orientation is defined as mod(*θ*_pref._, *π*). To determine the goodness of fit, we calculated the residual sum of squares of fitting error *E*, as well as the coefficient of determination *R*^2^:

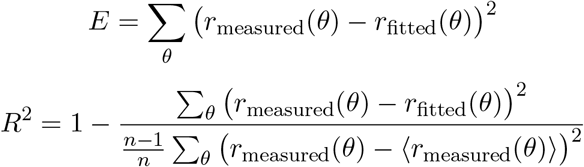

where *r*_measured_(*θ*) and *r*_fitted_(*θ*) are the experimental measured and fitted responses at direction *θ* respectively. ⟨·⟩ is the mean response across *θ*. Only when *E* < 0.4 and *R*^2^ > 0.6 the ROI was defined as well fit to the bimodal Gaussian function. And only when ROIs with tuning curves that were 1) responsive; 2) significant anisotropic across the 12 direction; 3) well fit to the bimodal Gaussian function, we define those ROIs as orientation selective (OS).

To quantify neuronal tuning we defined global orientation selectivity index (gOSI) for OS ROIs:

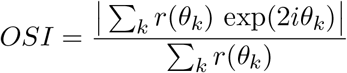

where k is the index among 12 directions, and *r*(*θ*) is the tuning curve. *gOSI* = 0 means no selectivity, and *gOSI* = 1 means perfect selectivity.

To calculate the distance dependent functions of tuning curve correlations in Figure 1-3, 5 and 7-8, we compute the cortical distance (defined as the distance between the center of mass of their cell bodies), and the correlation coefficient (i.e. normalized dot product) of tuning curves for all pairs of OS neurons. We then defined distance bins start from 5 *µm* with interval of 5 *µm*, i.e. 5-10 *µm*, 10-15 *µm*, etc.), and calculated the mean value and standard error of correlation values within all these bins for the error bar plots. Similar methods were used for the distance dependent functions of the difference between preferred orientation angles (Δ*θ*).

For trial-to-trial noise correlation measurements, the neuropil factor of *α* = 0.75 was used for all eight datasets, to avoid asymptotic correlation over broad distance that is possibly due to neuropil contamination. We determine whether ROIs are visually responsive by performing one-way ANOVA on the average Δ*F*/*F*_0_ during blank frames versus grating frames (*P* < 0.05). Out of visually responsive ROIs, we determine which ones are orientation selective by performing one-way ANOVA on the average Δ*F*/*F*_0_ during grating frames for each orientation separately (*P* < 0.05). Their preferred orientation is then defined as the orientation for which the average Δ*F*/*F*_0_ during grating frames is the largest. If there are pairs of ROIs which are closer than 5 *µm*, only the largest one of each pair is kept for the analyses. For noise correlations as a function of pairwise distance: we select only orientation selective ROIs and investigate only fluorescence during grating frames. For each grating orientation separately, we compute the Pearson correlation coefficient for the average response in the last 400 ms of each trial for each pair of selected ROIs. The results are grouped into the corresponding 5 *µm* pairwise distance bins and combined for all four grating orientations. In the condition with similar tuning, only ROIs whose preferred orientation is *θ* are selected for the *θ* grating frames, where *θ* ∈ {45, 135, 180, 270} (deg.).

### Single cell optogenetics

Holographic optogenetic experiments were performed as described previously (Oldenburg et al., 2024). Briefly: adult mice (age 6 - 12 weeks) of both sexes expressing GCaMP6s in excitatory neurons via tetO-GCaMP6s (Jackson Laboratory, 024742) × Camk2a-tTA (Jackson Laboratory, 003010) and Cre recombinase in a subset of excitatory neurons via SepW1-Cre (MGI:5519915). Mice were transfected with cre dependent ChroME2s (Sridharan et al., 2022), bicistronically linked to a nuclear localized mRuby3 used for targeting photostimulation, via Adeno-Associated Virus (AAV), (Addgene, 170163). In vivo mulitiphoton stimulation experiments were performed using a setup capable of 3D-SHOT, as described previously (Bounds et al., 2023; Mardinly et al., 2018; Oldenburg et al., 2023; Oldenburg et al., 2024; Pégard et al., 2017). The microscope is adapted on a movable objective microscope (Sutter Instrument) platform, with the addition of a temporally focused photostimulation path. Imaging was performed with a chameleon Ultra II (coherent), and photostimulation was performed with a Monaco40 (Coherent). Temporal focusing was achieved via a blazed holographic diffraction grating (Newport Corporation, R5000626767-19311), and holographic targeting via the high refresh rate spatial light modulator (SLM; HSP1920-1064-HSP8-HB, 1920 × 1152 pixels; Meadowlark Optics). To limit the risk of artifacts from photo stimulation, the stimulation laser was synchronized to the scan phase of the resonance galvos using an Arduino Mega (Arduino), gated to be only on the edges of every line scan.

Cells were targeted for stimulation based on the nuclear-localized mRuby3 signal bicistronically linked to the opsin. As described in Oldenburg et al., 2024, each putatively opsin positive cell was screened online for their ability to be stimulated with powers ranging from 12.5 to 100 mW per cell, to ensure both reliable activation and minimal power use. Each targeted cell was driven with ten 5 ms pulses at 10 Hz. Responding cells were analyzed offline using *Suite2p* (Pachitariu et al., 2017). Trials were excluded if the targeted cell failed to respond to photostimulation (at least 0.25 z-scored fluorescence above baseline), or if that target failed to respond in 50% of attempts. Cells were excluded from analysis if they were located above or below the targeted cell (within 22.5 *µm* radial), or 5 *µm* within the same plane.

### Spiking neuron network model

Our network of spatially organized spiking neuron models is build on previous models from our group (Gozel and Doiron, 2022; Huang et al., 2022; Huang et al., 2019; Rosenbaum and Doiron, 2014; Rosenbaum et al., 2017). To mimic the recorded L4 activity we build a model L4 with the same distance dependence of tuning curve correlation as in the data (Figure 2D). To this end, we first introduce an imaginary layer “X” (LX) that innervates L4. The LX population contains *N*_*X*_ = 10000 excitatory neurons arranged on uniform, two-dimensional squared grid (i.e. 100 × 100). We take the square domain to have length *L* = 750 *µm*, and the spacing between neighboring neurons is 7.5 *µm*. The firing rate of LX neuron *i* in response to oriented stimuli is given by a stimulus *θ* dependent mean corrupted by independent sources of noise:

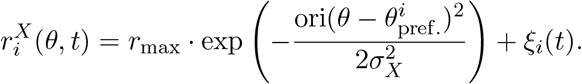

The stimulus orientation *θ* is taken from 10 orientations uniformly spaced between 0 and *π. r*_max_ = 10 Hz is the maximum firing rate and 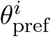 is the preferred orientation of neuron *i*. The organization of 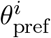 on the two-dimensional LX space is fully random, i.e. a strict “salt-and-pepper” map. The function ori(Δ*θ*) = min(|Δ*θ*|, |Δ*θ* − *π*|, |Δ*θ* + *π*|) wraps periodic orientation values onto the interval 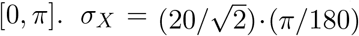 (rad) is the width of the tuning curve. The noise term *ξ*_*i*_(*t*) follows an Ornstein-Uhlenbeck (OU) process:

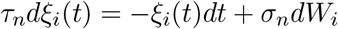

where *dt* = 1 (ms) in the simulation, τ_*n*_ = 40 (ms), *σ*_*n*_ = 0.03 (ms^−1*/*2^), and the increment of Wiener process *dW*_*i*_ ∼ 𝒩 (0, *σ*^2^ = *dt*). The Weiner processes are all independent of one another.

L4 contains *N*_*F*_ = 10000 excitatory Poisson-spiking neurons arranged on similar two-dimensional squared grids with the same length *L* = 750 *µm*, so that the interval between neighboring L4 neurons is 7.5 *µm*. Their firing rates are determined by:

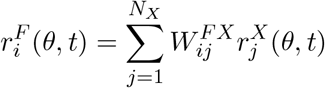

where 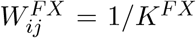 if LX neuron *j* connects to L4 neuron i, and 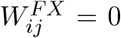 if not. *K*^*FX*^ = 60 is the expected number of connections that each L4 neuron receives from LX. The probability that a L4 neuron (with coordinate **x** = (*x*_1_, *x*_2_)) is post-synaptic to a LX neuron (with coordinate **y** = (*y*_1_, *y*_2_)) depends on their distance measured periodically on the two-dimensional square:

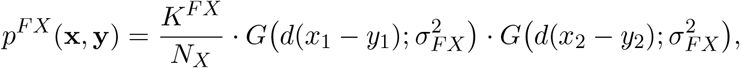

where 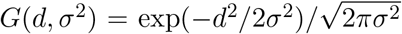 is a Gaussian-shaped function, and *d*(*x* − *y*) = min(|*x* − *y*|, |*x* − *y* − *L*|, |*x* − *y* + *L*|) wraps the distance onto the periodic interval [0, *L*]. We set *σ*_*FX*_ = 5.15 *µm* so that the firing rates of the L4 population has the same distance dependent tuning curve correlations as in the data (Figure 2D). Furthermore, the firing rates of all L4 neurons are scaled with a uniform factor so that the averaged peak firing rate of tuning curves is equal to *r*_max_ = 10 Hz. Spike trains of L4 neurons are then generated as inhomogeneous and independent Poisson processes with the rate 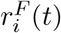.

The L2/3 contains recurrently coupled *N*_*E*_ = 10000 excitatory and *N*_*I*_ = 2500 inhibitory neurons arranged on similar two-dimensional squared grids with the same length *L* = 750 *µm*, so that the interval between neighboring E and I neurons are 7.5 *µm* and 15 *µm*, respectively. Each neuron is modeled as an exponential integrate-and-fire (EIF) neuron, and the membrane potential of neuron *i* in population *α* = *E, I* is determined by:

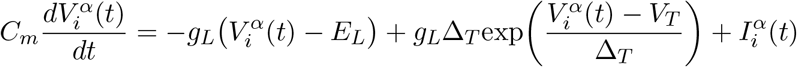

Once 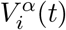 exceeds the threshold *V*_th_ the neuron spikes, and its membrane potential will be held for a refractory period τ_ref_ , and then reset to the value *V*_re_. For excitatory neurons, we take τ_*m*_ = *C*_*m*_/*g*_*L*_ = 15 ms, *E*_*L*_ = −60 mV, *V*_*T*_ = −50 mV, *V*_th_ = −10 mV, Δ_*T*_ = 2 mV, *V*_re_ = −65 mV and τ_ref_ = 1.5 ms. For inhibitory neurons, we take the same parameters except τ_*m*_ = 10 ms, Δ_*T*_ = 0.5 mV and τ_ref_ = 0.5 ms. The total input current to each neuron is:

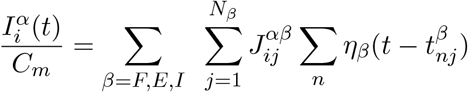

where β = *F, E, I* represents presynaptic neurons types (feedforward projection from L4, and *E* or *I* recurrent projection from L2/3); 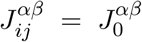 if neuron *j* from population β connects to neuron *i* in population *α*, and 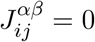 if not. The time point 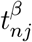 is the n^th^ spike time from neuron *j* in population β. The postsynaptic current function η_*β*_(Δ*t*) contains one fast and one slow component:

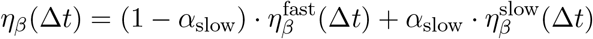

where

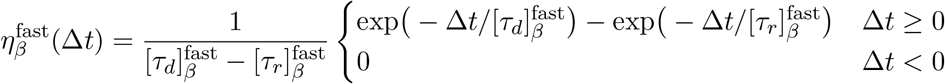

and 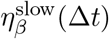 has similar form. We use *α*_slow_ = 0.3; [τ_*r*_]^fast^ = 1 ms for 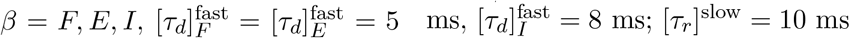 and [τ_*d*_]^slow^ = 100 ms for β = *F, E, I*.

The connection probability between a pair of neurons with coordinate **x** and **y** only depends on their distance:

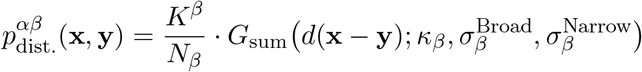

where *G*_sum_ is the sum of broad and narrow scale of Gaussian-shaped functions as in the Eq. (1), and *d*(**x** − **y**) calculates the distance between a pair of neuron on the periodic square [0, *L*] × [0, *L*]. Note that a postsynaptic neuron is allowed to have multiple synaptic connections to a single presynaptic neuron, so that some synaptic connections (especially for nearby connections) could have stronger postsynaptic potential than others. Also, we assume *κ, σ*^Broad^, *σ*^Narrow^ and the connection in-degree *K* depend on the presynaptic neuron type β only.

To compute tuning curves and tuning curve correlation of neurons in the network, we set up simulations containing alternative OFF (1000 ms) and ON (2000 ms) intervals. During OFF intervals, all L4 neurons fire spontaneously with uniform rate 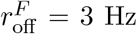. During ON intervals, each of 10 stimulus orientations that uniformly spaced between 0 and *π* are presented in LX for 100 trials (ON intervals), and spike counts during those ON intervals are used to compute the average response to each orientation, i.e. the tuning curve. The first OFF-ON interval (3000 ms) are discarded. The initial states of membrane potentials of each neuron are randomized in each simulation.

To create the final feedforward and recurrent wiring in our network we took a two staged approach. We first built a model where only the above spatial wiring was considered. We computed the tuning curves and preferred orientations of all L2/3 neurons. With these tuning measurements we then rewired all the feedforward and recurrent connections with new probability distribution that depends on both cortical distance, as well as the difference between preferred orientation angles of neurons, i.e. the ‘like-to-like’ feature-dependent rule in the Eq. (2) and (3):

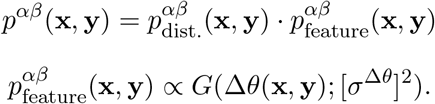

The final *p*^*αβ*^(**x, y**) should be properly normalized so that the sum equals to 1. We take *σ*^Δ*θ*^ = 40 · (*π*/180) (rad) for all types of feedforward and recurrent connections in the simulation.

After we generated samples of connections with the final joint probability distributions, we used maximum likelihood estimation to estimate the *κ, σ*^Broad^, *σ*^Narrow^ of the sum-of-Gaussians probability distribution (Eq. (1)). The results, summarized in the bottom panel (‘MODEL’ table) of Figure 6C, are different from the initial values we used, which could be due to the weak spatial structure in 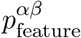 due to the micro-clustered organization, as well as the effect of finite interval distance between neighboring neurons. To compute trial-to-trial noise correlation, we use the same network parameters and connection matrices, and we choose 4 stimulus orientations that uniformly tiled between 0 and *π*, but each orientation was presented for 250 trials (ON intervals). The distance dependent function of noise correlation for each stimulus orientation were averaged to get the results in Figure 8B.

For the simulations of single cell perturbations in L2/3, we used the same network parameters and connection matrices, and we set up simulations containing alternative OFF (2000 ms) and ON (1000 ms) stimulation intervals. During both OFF and ON intervals, all L4 neurons fired spontaneously with a uniform rate 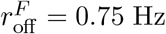. During ON intervals, one random neuron in L2/3 E population was forced to spike 10 times, with an interval of 100 ms. The spike counts of all the other neurons in the last 500 ms time bin of the ON interval were recorded. We simulated a total of 15000 trials (ON intervals) with different random L2/3 neuron stimulated to compute the averaged perturbed firing rates of L2/3 E population, as well as 50 trials without any stimulation to compute the basal firing rates. The distance dependent function of the change of average firing rate (ΔFR) are shown in Figure 7E-7F.

For all the spiking network simulations in the text (Figure 4*D* −4*G*, 5*C*_2_, 5*D*_2_, 5*E*_2_, 5*F*, 7*E* −7*F*, 8*B* and S7), we take 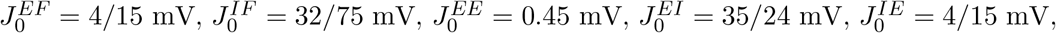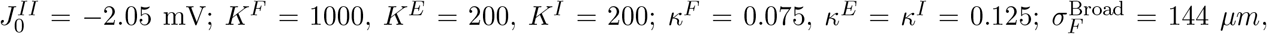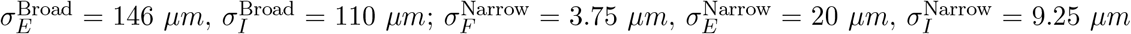. Note that the distance dependent function of tuning curve correlation and the orientation maps in Figure 5*C*_2_, 5*D*_2_ and 5*E*_2_ are generated with the spiking neuron network model, while the color maps in Figure 5*C*_1_, 5*D*_1_ and 5*E*_1_ are generated with a simplified linearized neural field model. The details of the linearized neural field model, and the way to compute the distance dependent function of L2/3 E tuning curve correlation with the assumption of a continuous of neural field (rather than neurons in grids) are introduced in the following Supplementary Methods section.

## Acknowledgments

B.D, N.J., H.A. and K.M are supported by the NIH grants U19NS107613-01. B.D. is supported by NIH grant R01NS133598 and CRCNS-R01EY034723, a Vannevar Bush faculty fellowship (N000141812002), and the Simons Foundation Collaboration on the Global Brain. This research benefited from Physics Frontier Center for Living Systems funded by the National Science Foundation (PHY-2317138). Financial support was provided via the National Institute from Mathematics and Theory in Biology (Simons Foundation award MP-TMPS-00005320 and NSF award DMS-2235451). H.A. is supported by NIH grant R01EY023756 and a Chan-Zuckerberg Biohub Investigatorship. K.M. is supported by NIH grants 5U01 NS108683 and 1R01 EY029999, and the Simons Foundation Collaboration on the Global Brain. M.D. is supported by a fellowship from the Kavli Foundation and NIH grants R21 EY035064. I.O. is supported by and NIH K99/R00EY029758, DP2MH140135, the Whitehall Foundation Award 2023-05-40, and the Searle Scholars Program Award SSP-2024-109. F.R. is funded by the Armenise-Harward Foundation (CDA to LFR) and the Human Technopole (HT–ECF 3588 to LFR).

## Supplementary Methods

In this supplementary method section we provide a linear approximation of responses of the *E* − *I* spiking neuron network. This allows us to explore the distance dependence of tuning correlations over a wide range of parameters (*σ*^Broad^, *σ*^Narrow^, *κ*).

### S1: Linearized neural field model

In this section we will derive the linearized model for Figure 5*C*_1_, 5*D*_1_ and 5*E*_1_.

Consider an *E* or *I* L2/3 neuron in the network (denoted by superscript *α* = *E, I*) with index *i* receiving a time-dependent input current 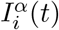. We decompose the total synaptic input current 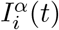 into the sum of feedforward (*F*), recurrent excitatory (*E*) and recurrent inhibitory (*I*) terms:

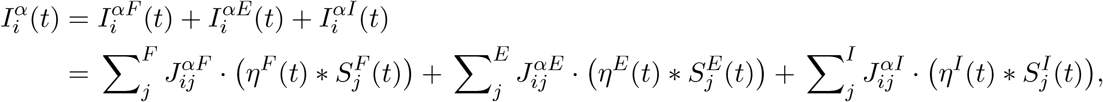

where η^*β*^(*t*) (β = *F, E, I*) is the synaptic current filter from population *β*, and 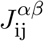 is the synaptic weight from neuron *j* in population *β* to neuron *i* in population *α*. The spike train response from neuron *j* in population *β* is 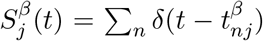, with 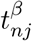 being the n^th^ spike time from neuron *j* in population *β*. The symbol ∗ denotes a temporal convolution: 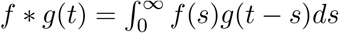.

Assuming weak input current (due to weak synaptic connections, for example) we approximate the neuronal response as:

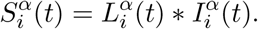

Here we have introduced the linear response kernel 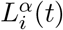, which is the filter that transfers input currents to output activity. We remark that the above equation is formally incorrect, since we cannot approximate 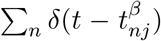 as a convolution of smooth, continuous functions. This approximation only makes sense after an expectation over realizations or time evolution of the stochastic terms *dW*_*i*_ from the LX inputs to L4 neurons (see Methods: Spiking neuron network model). We perform this expectation by considering equilibrium (statistical) activity and we take the temporal Fourier transform of the spike train 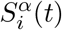,

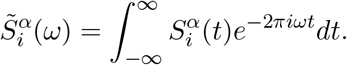

Then we have

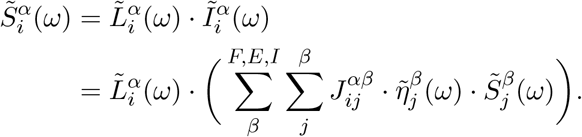

Since our study does not consider the dynamics of network activity, we then restrict our analysis to the zero frequency (ω = 0) case. Note that 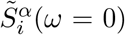 is the average spike count number, i.e. the mean firing rate of neuron, 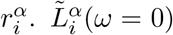 is simply the gain of the neuron, 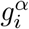. And 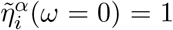 normalizes our synaptic current filter function. So in total we have

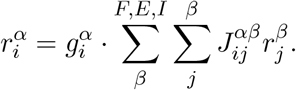

Let *W* 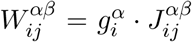 be the effective coupling, and using 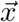 and 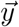 to represent the coordinate of neuron *i* and *j* on the two-dimensional plane gives

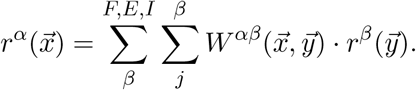

We assume that the effective connection weight 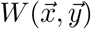 is translationally and circularly symmetric, i.e. it only depends on the distance between pre-synaptic and post-synaptic neurons (measured along the two-dimensional plane for L2/3 → L2/3 recurrent connections, and parallel to the plane for L4 → L2/3 feedforward connections), so that:

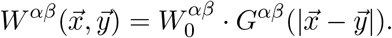

Here 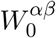 is the mean effective connection weight, and *G*(·) is an arbitrary function defining the shape of connectivity. It could be a Gaussian-shaped function, or the sum of broad-and-narrow Gaussians in the later sections. So, 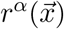 becomes:

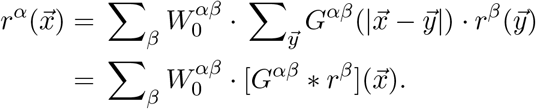

Consider the spatial Fourier transform of 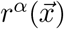:

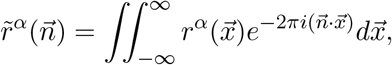

giving:

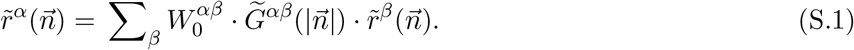

We rewrite Eq. (S.1) in the form of 2 × 2 matrices (one for each 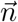):

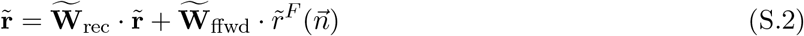

where

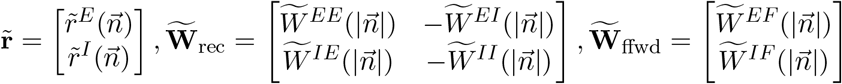

and

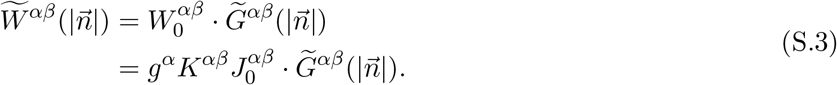

Here *g*^*α*^ is the gain of a neuron in population *α, K*^*αβ*^ is the number of connections from population *β* to *α*, and 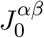 is the post-synaptic potential of an individual synapses from *β* to *α*. Finally, 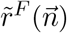 is the spatial organization of activity in the feedforward layer L4.

We solve Eq. (S.2) to get the spatial organization of firing rates of L2/3 *E* and *I* populations in the Fourier space:

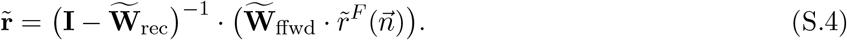

To compute the tuning curve correlations of neuron pairs over distance (defined as the normalized dot product of tuning curves), it is equivalent to consider the covariance function of the spatial profile of firing rates averaged over stimuli orientations:

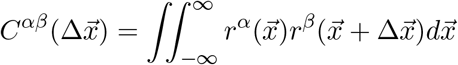

According to the Wiener-Khinchin theorem, the covariance function in the Fourier space is equal to the absolute square of 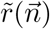:

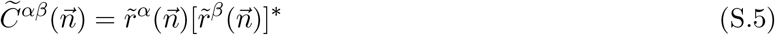

So we have

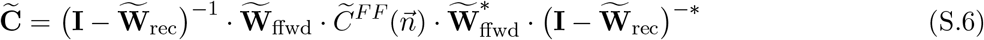

where

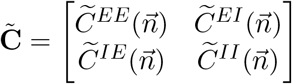

and 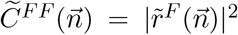 is the Fourier transform of the correlation structure in the feedforward layer L4, which is enforced and organized to be the same as in data (Figure 2D). Thus, with the knowledge of neural gain *g*^*α*^, connection number (or equivalent probability) *K*^*αβ*^, postsynaptic potential 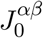, as well as the spatial profile *G*^*αβ*^(*d*) of feedforward and recurrent connections, we can calculate the output signal correlation function over distance for the L2/3 E or I populations. The connection number *K* and postsynaptic potential *J*_0_ are set in the Methods. To determine the neural gain *g*^*E*^ and *g*^*I*^, we sampled the synaptic input currents (*I*) and firing rate *r* of 750 excitatory and 750 inhibitory neurons, and fit the *r* and *I* relationship with the following threshold power function:

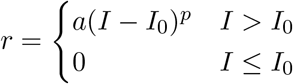

Once we obtain the best fit a, *I*_0_ and *p*, we determine the approximated gain with the derivative of the *r*(*I*) function at the point where *r* is equal to the mean firing rate of the *E* of *I* populations, i.e. at their operating points.

To derive the distance dependent function of trial-to-trial noise correlation between L2/3 spike trains we have:

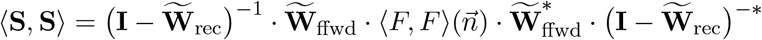

where

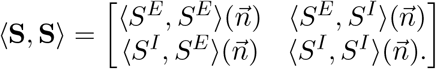

We note that the trial-to-trial noise covariance in L4, ⟨*F, F*⟩(*d*), is proportional to a two-dimensional Dirac delta function, since the dynamics of L4 neuron in the model follows independent OU process (see Methods), and different neurons have zero trial-to-trial correlation. So in the Fourier space 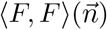 should be a constant *C*_0_**I**, with *C*_0_ a constant. This yields:

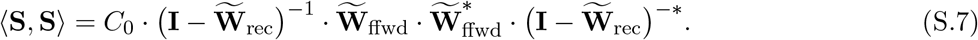

implying that the trial-to-trial noise covariance in L2/3 is a reflection of both the feedforward and recurrent wiring structure.

To generate the color maps in Figure 5*C*_1_, 5*D*_1_ and 5*E*_1_, for terms 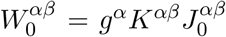 in the neural field model (Eq. S.3), we take the same parameters *K*^*αβ*^ and 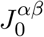 as in the ‘Spiking neuron networks model’ part of the Methods section, and we take *g*^*E*^ = 12 (*V* ^−1^) and *g*^*I*^ = 15 (*V* ^−1^). In Figure 5*C*_1_, we fix all *κ* = 0. In Figure 5*D*_1_, we fix *κ*^*F*^ = 0.075, *κ*^*E*^ = *κ*^*I*^ = 0.125, and 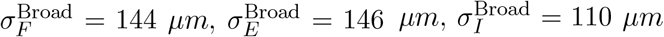. In Figure 5*E*_1_, we fix 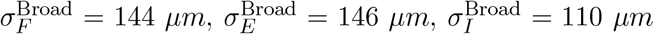, and 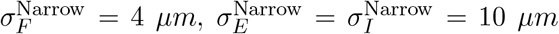. In Figure 5*D*_1_ and 5*E*_1_, the saturation and hue of each point on the color map represents the normalized amplitude and the standard deviation of Gaussian fits to the distance dependent function of L2/3 E tuning curve correlation. In Figure 5*C*_1_, since the correlation function is no longer monotonically decreasing as a Gaussian-shaped function, we define the spatial scale λ_0_ of a non-monotonic function as the inverse of the spatial mode with maximum response in the spatial Fourier space.

### S2: Conditions for spatially balanced solutions with Gaussian-shaped connectivity in the continuum limit

Following past studies (Rosenbaum and Doiron, 2014; Rosenbaum et al., 2017), in this section we will derive the conditions on the existence of a balanced solution over all spatial locations, i.e. Eq. (8) and (9) in the main text, in the framework of a linearized E-I model. According to Eq. (S.1) and Eq. (S.3), the firing rate of population *α* in Fourier space is

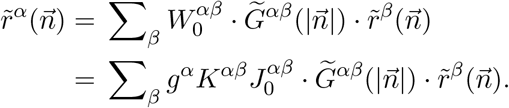

Following previous studies in balanced etworks (van Vreeswijk and Sompolinsky, 1996, 1998), we have the postsynaptic potential 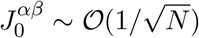 (where N is the total neuron number of the network) in order to maintain a *𝒪*(1) response correlation. Since *K*^*αβ*^ ∼ *𝒪*(*N*), we define

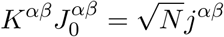

so that *j*^*αβ*^ ∼ *𝒪*(1):

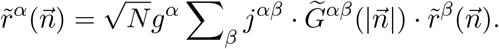

To maintain *r*^*α*^ ∼ *𝒪*(1), we must have that

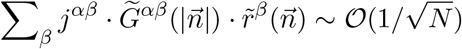

over all spatial Fourier modes 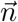. In the limit of *N* → ∞ (i.e. continuum limit), this means

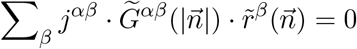

Rewrite this as a 2 × 2 matrix in the form:

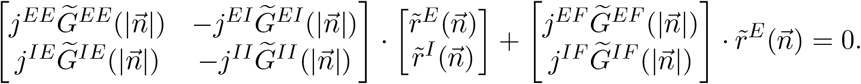

The spatial profiles of feedforward and recurrent connections, *G*^*αβ*^(*d*), are a normalized two-dimensional Gaussian-shaped function:

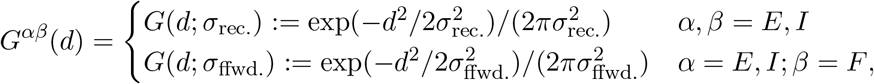

where *σ*_ffwd._ and *σ*_rec._ are the width (standard deviation) for feedforward and recurrent connections – we assume equal *σ*_ffwd._ for *G*^*EF*^ and *G*^*IF*^, and equal *σ*_rec._ for *G*^*EE*^, *G*^*EI*^, *G*^*IE*^ and *G*^*II*^. In the Fourier domain, those Gaussian-shaped functions are given by

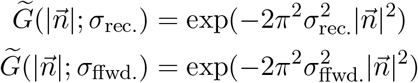

Thus, a balanced solution over all spatial Fourier modes requires:

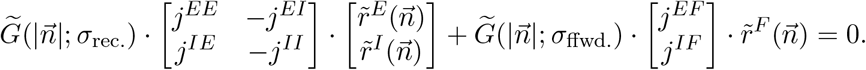

Solving this equation yields:

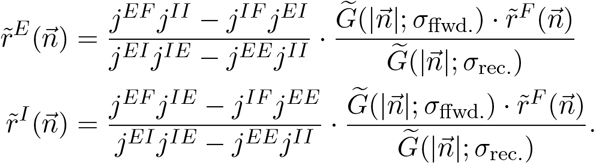

Thus, the spatial profile of both 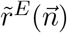 and 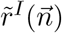 are determined by the factor

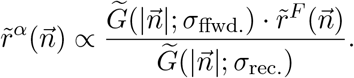

According to the Wiener-Khinchin theorem (Eq. S.5), 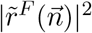 is equal to the auto-covariance function of firing rate 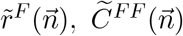 – the tuning curve correlation as a function of distance (spatial Fourier modes) in L4 (the feedforward layer). Thus, the standard deviation of Gaussian-shaped 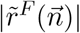, denoted as 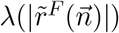, should be 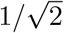 times of that of the distance dependent function of tuning correlation in L4 (∼ 9.16 *µm*, Figure 2D of the main text).

So, we have

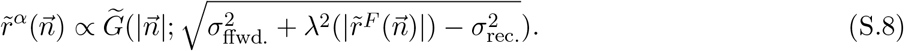

A valid real solution of Eq. (S.8) requires:

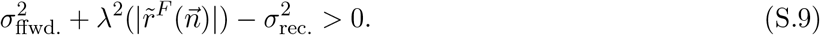

The inverse Fourier transform of 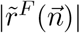 is the Δ*r*_*L*4_(*d*) we defined in the main text: the average deviation of firing rate of all neurons at a distance *d* relative to a reference L4 neuron, then averaged over all possible reference neurons. Thus, Eq. (S.9) is exactly the same as Eq. (8) of the main text which operates in the physical space 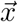 as opposed to Fourier space 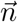. Also, the spatial scale in Eq. (S.8) is the same as λ(Δ*r*_*L*2*/*3_) in the Eq. (7) of the main text. Violating this condition will lead to a failure of the balanced solution at all spatial Fourier modes, and the network will instead yield a localized spatial pattern.

### S3: Linearized neural field model for single cell optogenetics

In this section we provide a theory to predict the spatial profile of neural responses to single cell optogenetic activation (Figure 7E-7F of the main text), based on the linearized E-I neural field model. We start from Eq. (S.2):

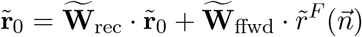

where **r**_0_ is the equilibrium state before optogenetic stimulation. The single cell optogenetic activation forces the target neuron to fire, and affects the firing rate of other neurons through recurrent wiring. We model such a perturbation through the multiplicative term **W**_rec_ · *δ***r**_*p*_:

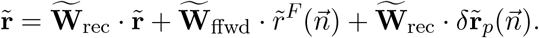

Taking the difference of the two equations and defining the firing rate change due to the perturbation as Δ**r** = **r** − **r**_0_, we have:

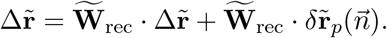

Solving for 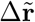 gives:

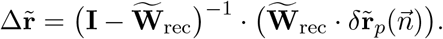

We define 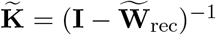 and write:

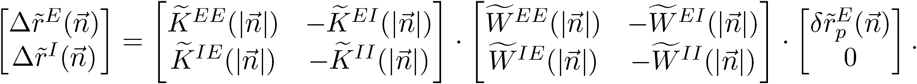

Note that the perturbation targets only a L2/3 *E* neuron, so 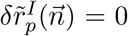. For the *E* population, we then have:

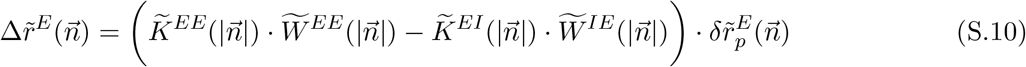

Here 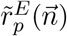 is the Fourier transform of a two-dimensional Dirac delta function, since only one single L2/3 E neuron is activated:

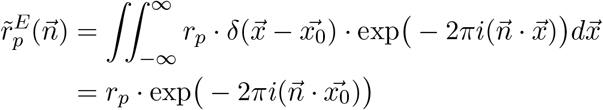

where 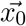 is the coordinate of the activated neuron, and *r*_*p*_ is the activation firing rate. Now we denote 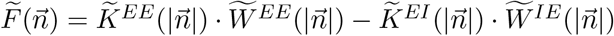, and take the inverse Fourier transform of Eq. (S.10):

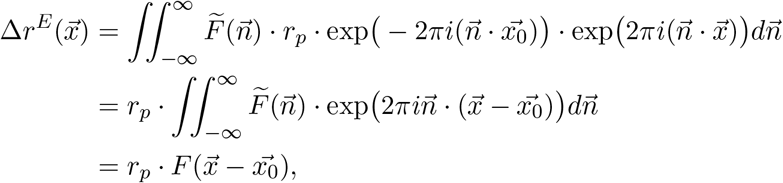

and

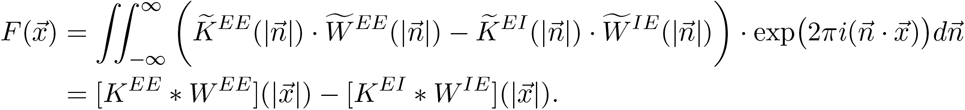

Thus, we have

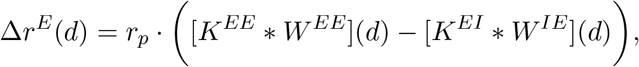

where 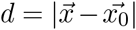 is the cortical distance between a neuron at position 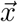 and the optogenetically activated neuron at position 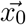.

Since the single neuron activation only provides a weak perturbation to the network, we approximate **K** = (**I** − **W**_rec_)^−1^ with the first-order expansion **K** ≈ **I** + **W**_rec_, so that

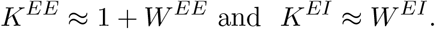

So finally the distance dependent function of the firing rate difference due to the single *E* neuron activation becomes:

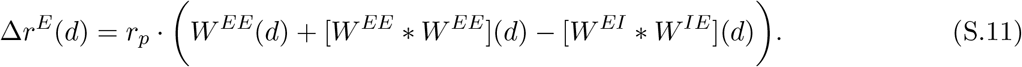

The three terms in Eq. (S.11), *W*^*EE*^(*d*), [*W*^*EE*^ ∗ *W*^*EE*^](*d*) and −[*W*^*EI*^ ∗ *W*^*IE*^](*d*), represent the effect of the *E* → *E* monosynaptic recurrent circuitry, the *E* → *E* → *E* and *E* → *I* → *E* disynaptic recurrent circuitry, respectively.

Let the spatial profile of *W*^*EE*^(*d*), *W*^*EI*^(*d*) and *W*^*IE*^(*d*) be the sum of one narrow and one broad Gaussian-shaped function:

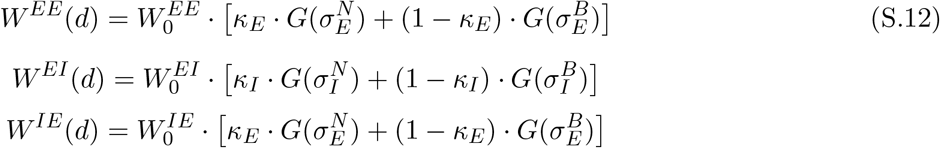

where *G*(*σ*) is normalized two-dimensional Gaussian-shaped function exp(−*d*^2^/2*σ*^2^)/(2*πσ*^2^); the superscript {*N, B*} indicates the narrow / broad component, and the subscript {*E, I*} indicates connection projected from *E* or *I* neuron; the effective connection weight is 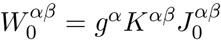, as in Eq. (S.3).

For the term [*W*^*EE*^ ∗ *W*^*EE*^](*d*) in Eq. (S.11), we have that

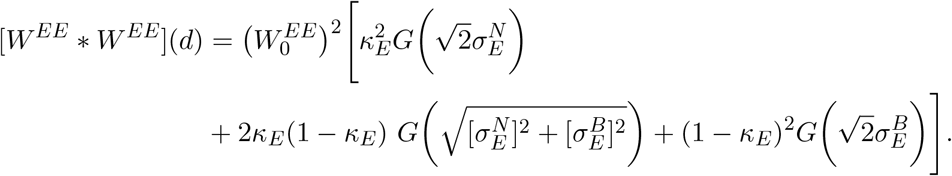

We note that 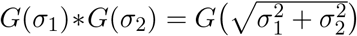. We make two more approximations: 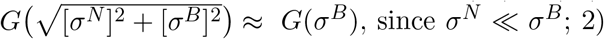, since *σN* ≪ *σB*;2) ignore any *𝒪*(*κ*^2^) terms, as well as any *𝒪*(*κ*) terms with broad spatial scales, since both have much weaker amplitudes than *𝒪*(*κ*) terms with narrow spatial scales in the term *W*^*EE*^(*d*) (Eq. S.12). With these approximations we have:

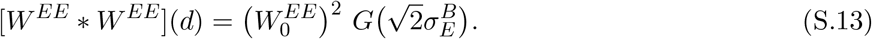

Similarly for the term [*W*^*EI*^ ∗ *W*^*IE*^ ](*d*) we have:

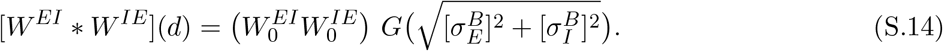

Inserting Eqs. (S.12), (S.13) and (S.14) into Eq. (S.11), we get the final result

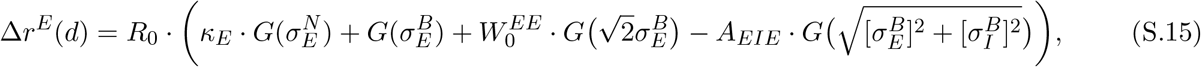

where 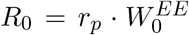, and 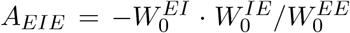. Note that the only narrow component term, 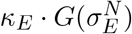, comes from the *E* → *E* monosynaptic interaction term *W*^*EE*^(*d*); the disynaptic *E* → *E* → *E* term [*W*^*EE*^ ∗ *W*^*EE*^](*d*) and *E* → *I* → *E* term [*W*^*EI*^ ∗ *W*^*IE*^](*d*) affect only the broad spatial scale of the result.

For the curves in Figure 7E and Figure 7F, we take *R*_0_ = 210 (Hz); for 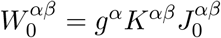, we take the same parameters *K*^*αβ*^ and 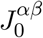 as in the simulation of the spiking neural network, and *g*^*E*^ = 12 (*V* ^−1^) and *g*^*I*^ = 15 (*V* ^−1^).

## Supplemental Information

**Supplementary Table 1:**
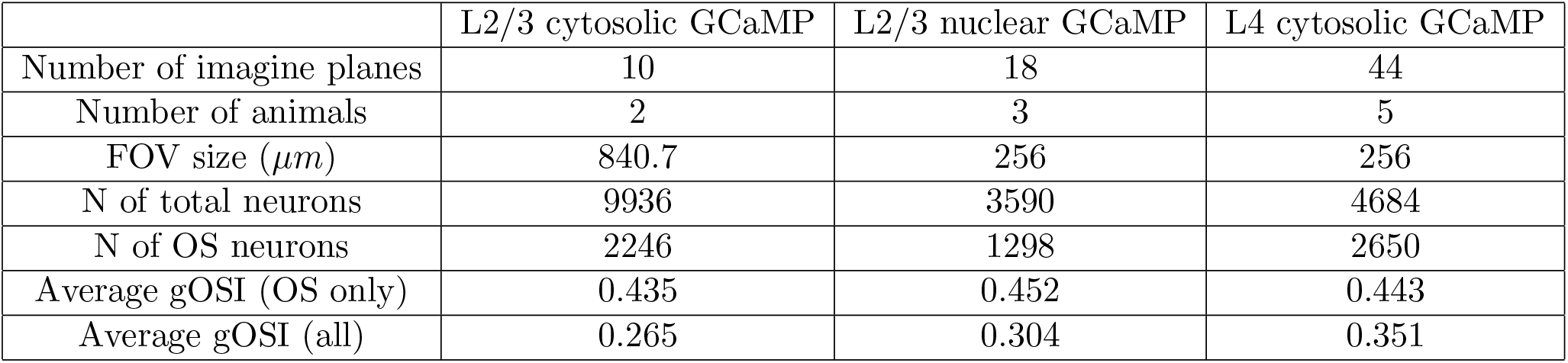
(related to Figure 1 and 2) Parameters of cytosolic GCaMP expressing L2/3, nuclear-targeted GCaMP expressing L2/3, and cytosolic GCaMP expressing L4 datasets.

|  | L2/3 cytosolic GCaMP | L2/3 nuclear GCaMP | L4 cytosolic GCaMP |
| --- | --- | --- | --- |
| Number of image planes | 10 | 18 | 44 |
| Number of animals | 2 | 3 | 5 |
| FOV size ($\mu m$) | 840.7 | 256 | 256 |
| N of total neurons | 9936 | 3590 | 4684 |
| N of OS neurons | 2246 | 1298 | 2650 |
| Average gOSI (OS only) | 0.435 | 0.452 | 0.443 |
| Average gOSI (all) | 0.265 | 0.304 | 0.351 |

**Supplementary Figure 1:**
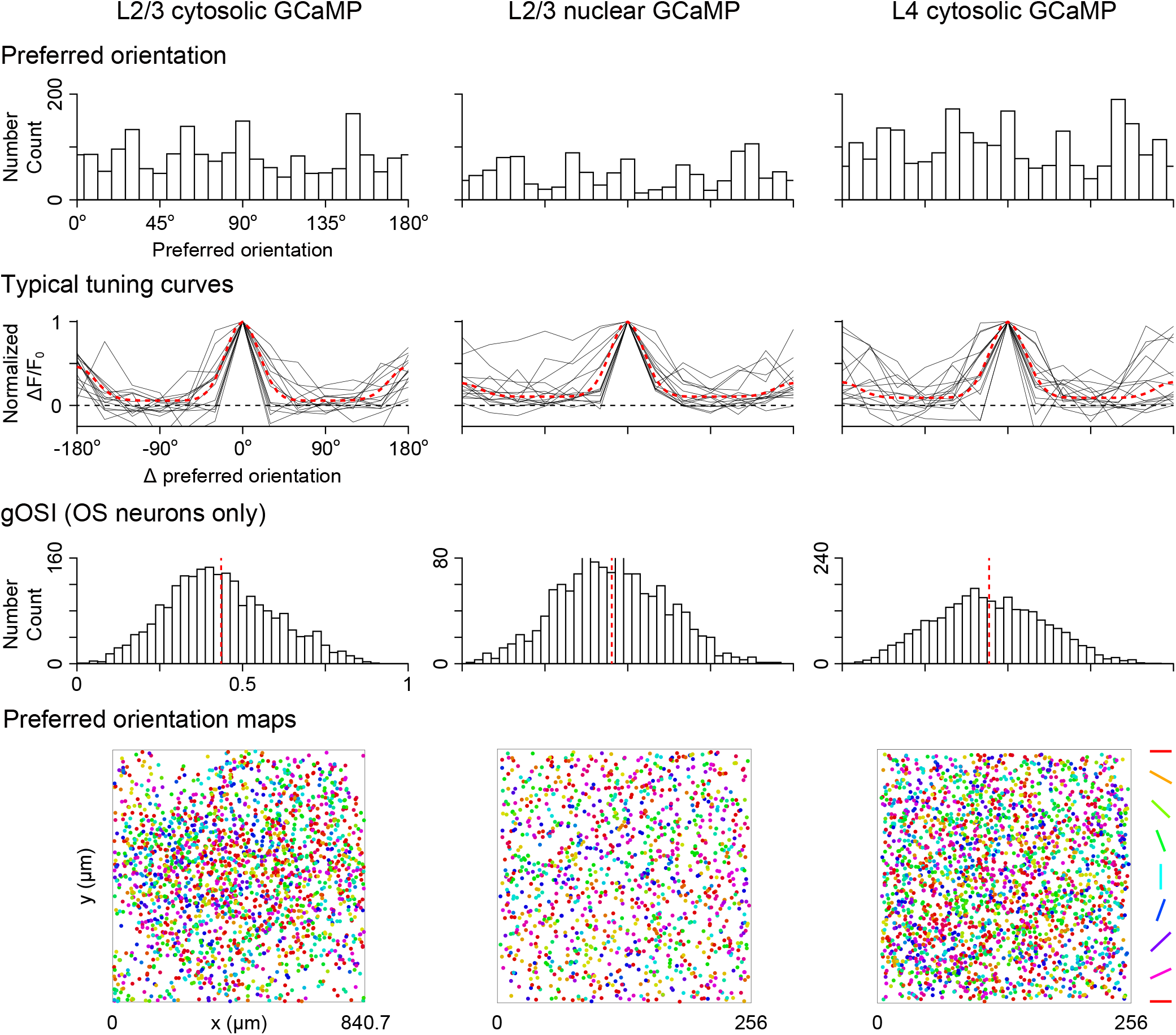
(related to Figure 1 and 2) The distribution of the preferred orientation, the normalized tuning curves of 15 random individual examples (gray) as well as the population average (red dashed) relative to preferred directions, the distribution of global orientation selectivity index (gOSI), and the preferred orientation map (vertically stacked), of all OS neurons in cytosolic GCaMP expressing L2/3, nuclear-targeted GCaMP expressing L2/3, and cytosolic GCaMP expressing L4 datasets.

**Supplementary Figure 2:**
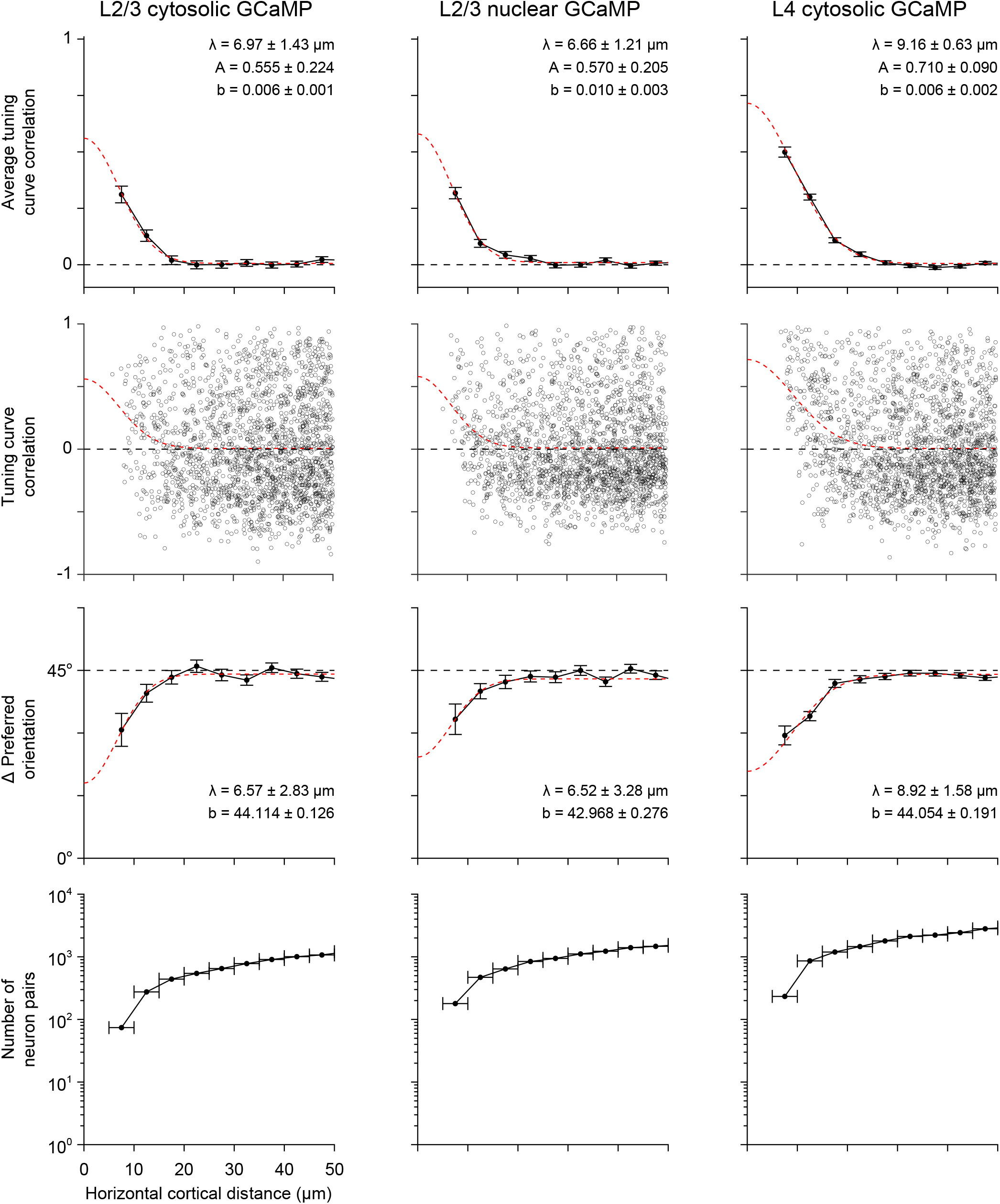
(related to Figure 1 and 2) For cytosolic GCaMP expressing L2/3, nuclear-targeted GCaMP expressing L2/3, and cytosolic GCaMP expressing L4 datasets, Row 1: The average tuning curve correlation in each distance bin; Row 2: Scatters of tuning curve correlation versus distance. Red dashed curve: same as in the Row 1; Row 3: The average difference of preferred orientations in each distance bin; Row 4: Numbers of pairs of neurons in each distance bin.

**Supplementary Figure 3:**
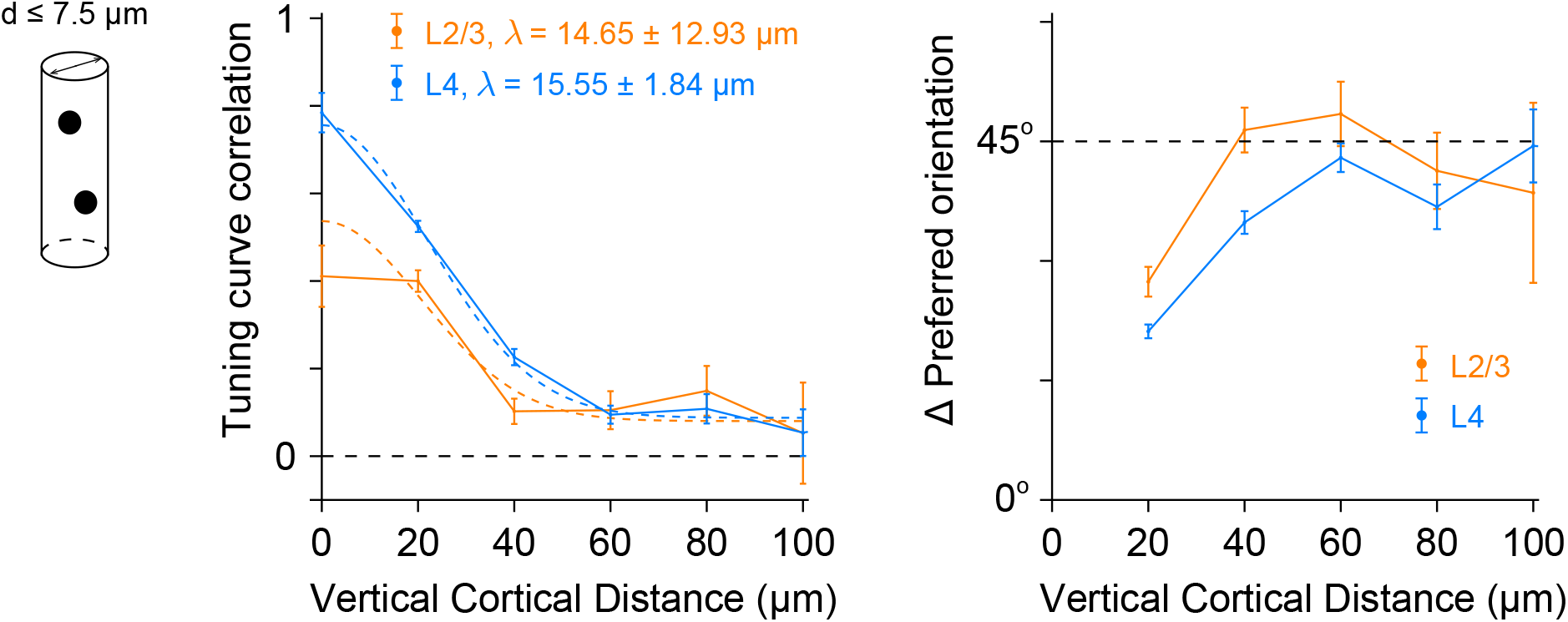
(related to Figure 1 and 2) The averaged tuning curve correlation and the difference of preferred orientations over vertical cortical distance, for neuron pairs whose horizontal cortical distance was less than 7.5 *µm*. Orange curve: L2/3 nuclear GCaMP; Blue curve: L4 cytosolic GCaMP.

**Supplementary Figure 4:**
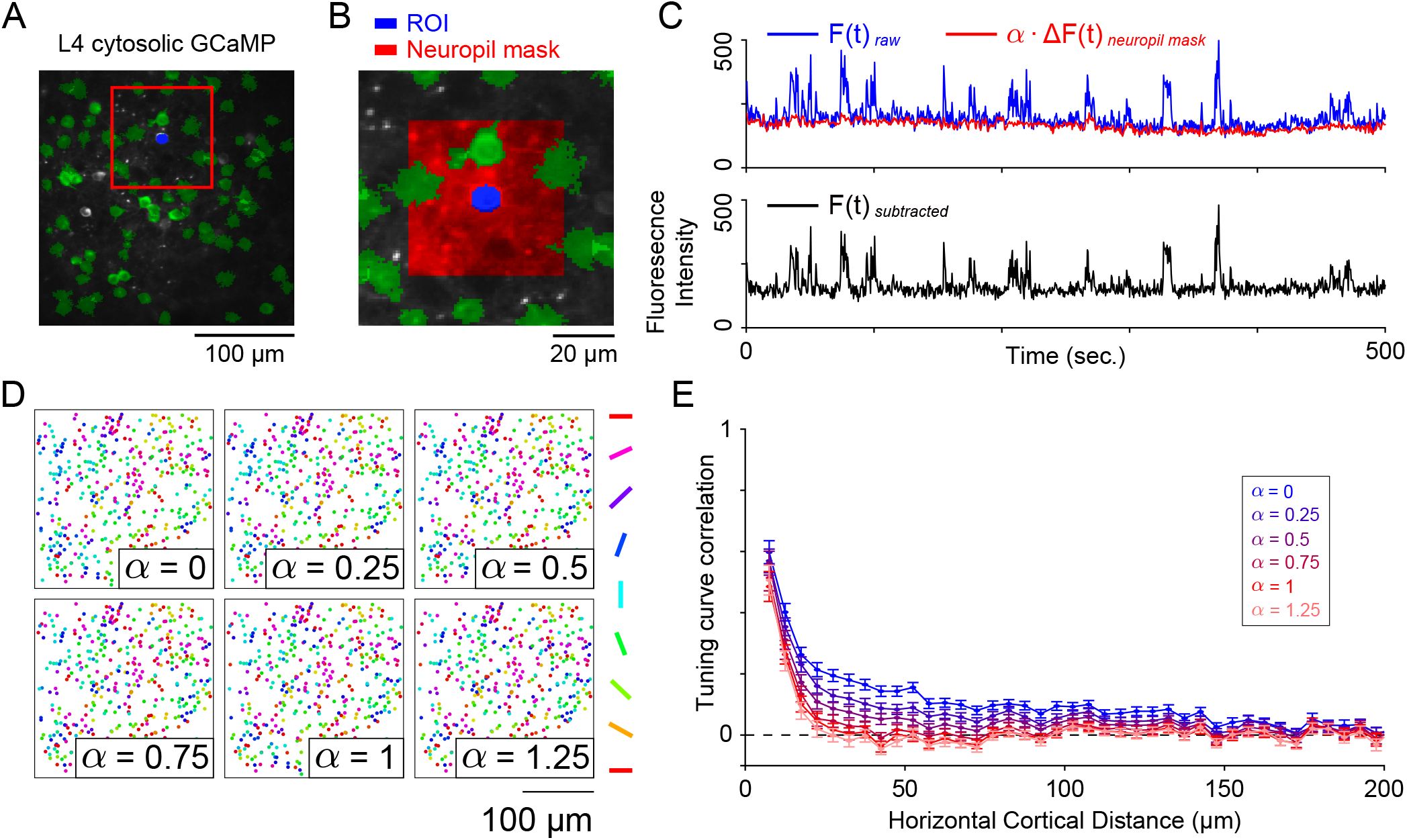
(related to Figure 3) Measuring the effect of neuropil subtraction on micro-clustered organization in cytosolic expressing L4 dataset. (A-E) organized as in Figure 3A, 3B, 3C, 3G and 3H.

**Supplementary Figure 5:**
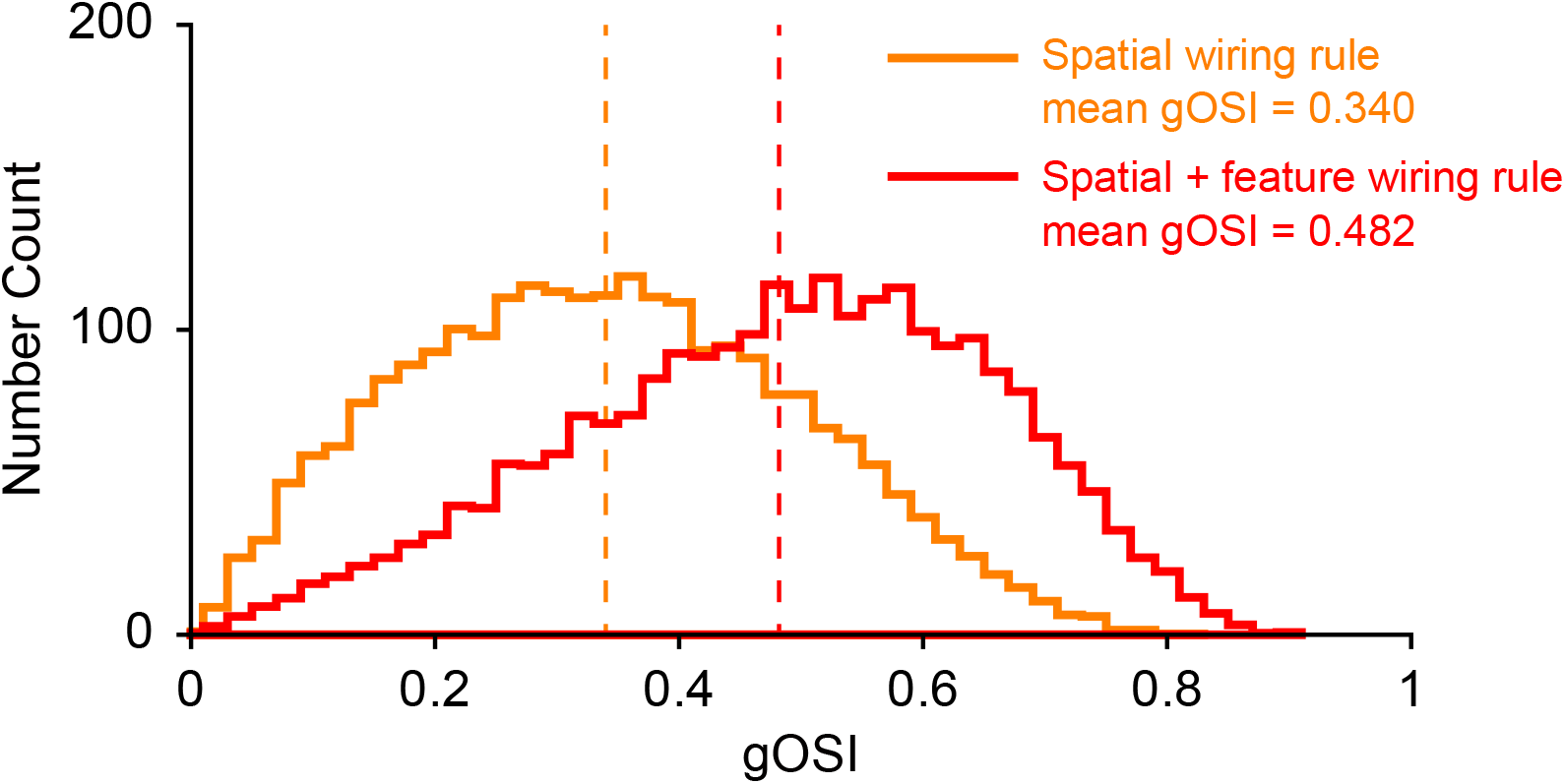
The orientation selectivity index of L2/3 E population for the network model where connection probability depends on distance only, or considers both distance and the ‘like-to-like’ feature dependent rule. Dashed lines represent the mean values. The latter one shows better orientation selectivity, while the model without feature dependent rule also show significant orientation selectivity.

**Supplementary Figure 6:**
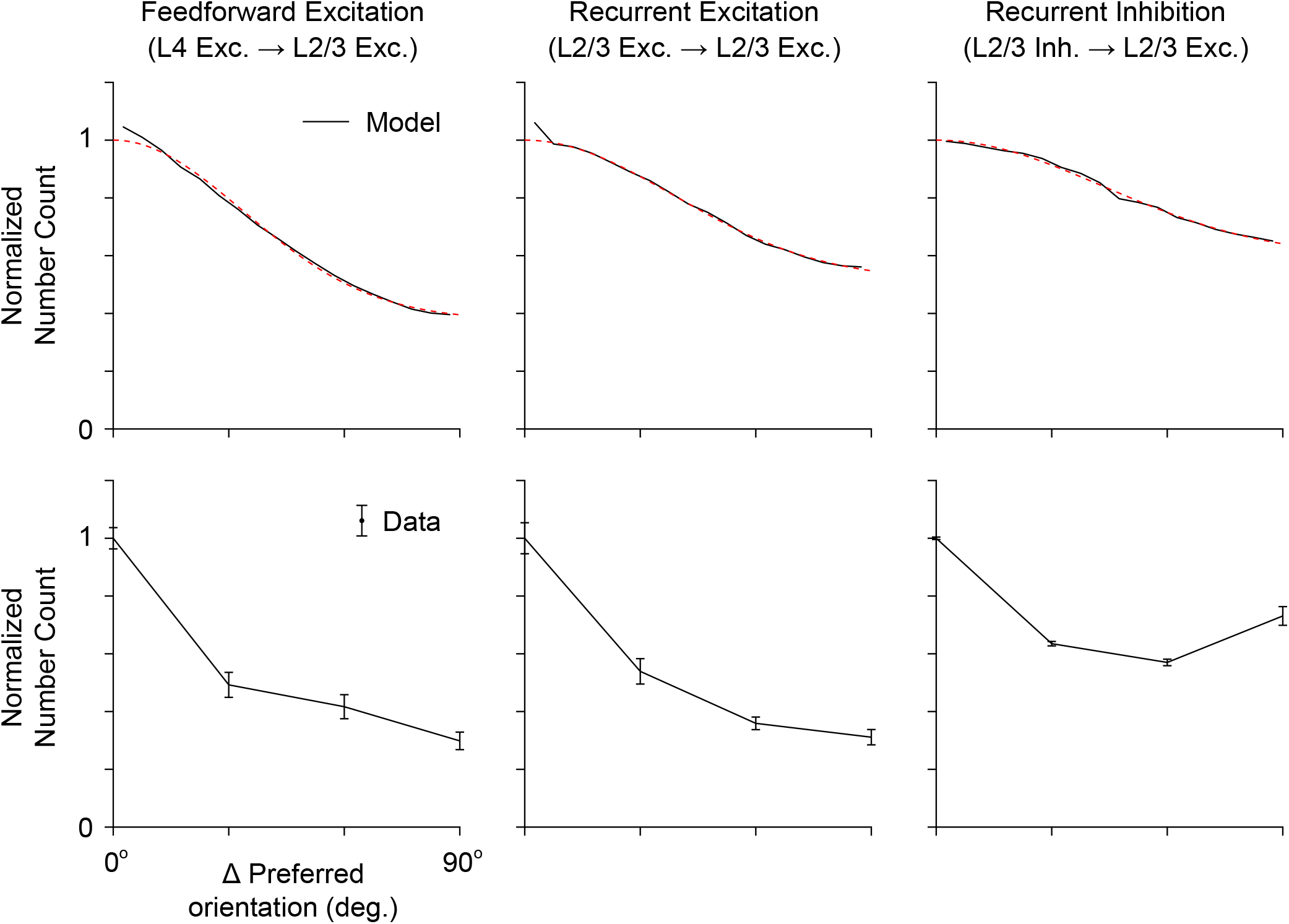
(related to Figure 6) The ‘like-to-like’ feature (Δ preferred orientation) dependent wiring rule in the network model (Black solid line: the network model; Red dashed line: Gaussian fit), and in the rabies tracing dataset (Black error-bars).

**Supplementary Figure 7:**
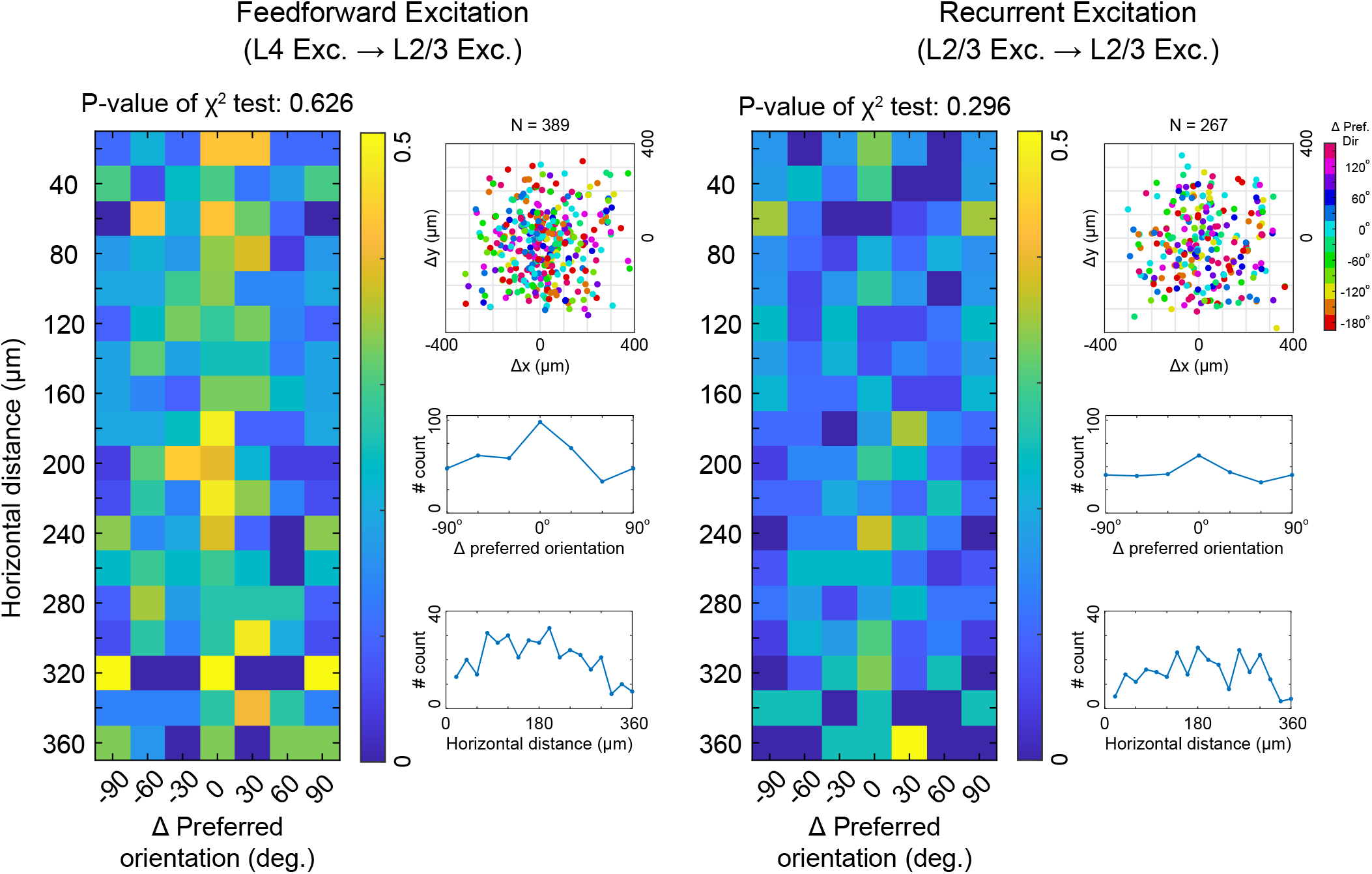
(related to Figure 6) The Chi-square test of independence for the spatial-dependent term and the feature-dependent term for the feedforward and excitatory recurrent connections in the rabies tracing dataset. Left: Two-dimensional joint probability distribution of connections over horizontal cortical distance, as well as the difference of preferred orientations. Color bar: number count normalized by the maximum value of each row. The p-values of Chi-square test are 0.626 and 0.296 for the feedforward and excitatory recurrent connections. Thus, we could not reject the null hypothesis of independence. Top right: Relative coordinates (Δ*x*, Δ*y*) of presynaptic ensembles stacked vertically (over z). The origin represents the location of postsynaptic L2/3 pyramidal neurons (n = 15, overlapped). Color represents the difference of preferred orientation between the corresponding postsynaptic neuron. Boundary between L2/3 and L4: 370 *µm* below the pia. Middle and bottom right: Marginal distributions on horizontal cortical distance, as well as on difference of preferred orientations.

## Notes

### Competing Interest Statement

The authors have declared no competing interest.

